# Characterization of a novel mesophilic CTP-dependent riboflavin kinase and rational engineering to create its thermostable homologs

**DOI:** 10.1101/2021.04.15.440061

**Authors:** Yashwant Kumar, Reman Kumar Singh, Amrita Brajagopal Hazra

## Abstract

Flavins play a central role in cellular metabolism as molecules that catalyze a wide range of oxidation-reduction reactions in living organisms. Several interesting variations in flavin biosynthesis exist among the domains of life, and the analysis of enzymes on this pathway have put forth many unique structural and mechanistic insights till date. The CTP-dependent riboflavin kinase in archaea is one such example - unlike most kinase enzymes that use adenosine triphosphate to conduct phosphorylation reactions, riboflavin kinases from archaea utilizes cytidine triphosphate (CTP) to phosphorylate riboflavin to produce flavin mononucleotide (FMN). In this study, we present the characterization of a new mesophilic archaeal riboflavin kinase homolog from *Methanococcus maripaludis* (*Mmp*RibK), which is linked closely in sequence to the previously characterized thermophilic homolog from *Methanocaldococcus jannaschii* (*Mj*RibK). We reconstitute the activity of the CTP-dependent *Mmp*RibK, determine its kinetic parameters, and analyse the molecular factors that contribute to the uncommon properties of this class of enzymes. Specifically, we probe the flexibility of *Mmp*RibK and *Mj*RibK under varying temperatures and the role of a metal ion for substrate binding and catalysis using molecular dynamics simulation and a series of experiments. Furthermore, based on the high degree of sequence similarity between the mesophilic *Mmp*RibK and the thermophilic *Mj*RibK, we use comparative analysis and site-directed mutagenesis to establish a set of the residues that are responsible for the thermostability of the enzyme without any loss in activity or substrate specificity. Our work contributes to the molecular understanding of flavin biosynthesis in archaea through the characterization of the first mesophilic CTP-dependent riboflavin kinase. Finally, it validates the role of salt bridges and rigidifying amino acid residues in imparting thermostability to enzymes, with implications in enzyme engineering and biotechnological applications.

## INTRODUCTION

Flavin cofactors such as flavin mononucleoide (FMN) and flavin adenine dinucleotide (FAD) are essential for primary metabolism in living organisms. The biosynthesis of flavins includes several steps that are mechanistically intriguing. Especially in archaea, the enzymes involved in the flavin biosynthesis pathway differ significantly in their sequence, structure, and mechanism as compared to their bacterial and eukaryotic counterparts.^[1–6]^ For example, the first enzyme GTP cyclohydrolase in archaea carries out a ring-opening hydrolysis on the purine imidazole ring of GTP to produce a formlyated intermediate unique to archaea, which is then converted to the diaminopyrimidine product by a formamide hydrolase enzyme.^[3,4]^ Another unusual example is the archaeal riboflavin synthase that catalyzes a dismutation reaction with two molecules of 6,7-dimethyl-8-ribityllumazine in the step leading to the formation of riboflavin. The pentacyclic adduct intermediate formed by the archaeal riboflavin synthase is a diastereomer of the corresponding bacterial adduct.^[6]^ The example that we analyse in this study is the archaeal riboflavin kinase, which converts riboflavin (Vitamin B_2_) into FMN via a phosphorylation reaction. This class of enzymes is unique in several ways: (a) in most known homologs of riboflavin kinase, this conversion is effected by the use of ATP. However, the riboflavin kinase homolog from the thermophilic archaea *Methanocaldococcus janaschii* (*Mj*RibK), the first characterized enzyme in this class, uses CTP as a source of phosphate, and is specific for its utilization.^[2,7]^ The only perceptible difference between the various nucleotide triphosphates is the nucleobase, thus the structural fold and mechanism of this enzyme appear to have adapted to selecting cytosine over other nucleobases. (b) The characterization of *Mj*RibK showed that the enzyme is active at 70°C without using a divalent metal ion, which was intriguing. Divalent metal cations such as Mg^2+^, Mn^2+^ and Zn^2+^ play an important role of stabilizing the negatively charged triphosphate group of the nucleotide and assisting in catalysis, with metal-independent behaviour only being exhibited by a handful of kinases/ pseudokinases.^[8–10]^ (c*) Mj*RibK has a sequence and fold that is unique with no similarity to any other known class of kinase, including a lack of the conventional phosphate binding motif in its active site.^[7]^ Such curious features make this class of CTP-dependent riboflavin kinases uniquely distinct from other known kinases and lend themselves to mechanistic investigations and enzyme engineering applications. However, as *Mj*RibK shows optimal activity at higher temperatures, it is not well-suited to extensive mechanistic explorations and has limited scope under physiological conditions. As the only reported CTP-dependent riboflavin kinases in literature are either from hyperthermophiles or thermophiles, we launched a search for a suitable mesophilic candidate that would be amenable to our studies.^[2,5,7]^ Bioinformatic and phylogeny analysis revealed that *Methanococcus maripaludis*, a closely related mesophilic archaea found in salt marshes, contains a homolog of the CTP-dependent riboflavin kinase (*Mmp*RibK). In this study, we report the characterization of *Mmp*RibK. We explore the functional differences between the *Mmp*RibK and *Mj*RibK homologs using biochemical tools and molecular modelling and establish the reason for several of the features that are characteristic of this class of enzymes. Further, we identify key amino acid residues responsible for the difference in activity of *Mj*RibK and *Mmp*RibK at different temperatures and create mutants of *Mmp*RibK which can operate up to 15°C higher than the wild-type enzyme without loss of activity or substrate specificity. Our findings explain some of the unique properties of the archaeal CTP-dependent riboflavin kinases and establish the role of salt bridges made by oppositely charged residue pairs such as Glu-Lys, Asp-Arg and Asp-Lys and residues that confer rigidity (such as prolines) in creating enzymes that can operate at higher temperatures.

## RESULTS AND DISCUSSION

### Bioinformatic and phylogenetic analyses reveal that *MmpRibK* is a close mesophilic homologue of the thermophilic enzyme *Mj*RibK

To identify a suitable mesophilic homolog for our study, we conducted a protein BLAST analysis with the sequence of the archaeal riboflavin kinase enzyme *Mj*RibK. The results revealed 1813 archaeal riboflavin kinase enzymes (EC: 2.7.1.161). Among the top 25 conserved sequences, 44% are mesophilic of which 32% are different strains of *M. maripludis* (Table S1). Next, we curated a set of 56 archaeal riboflavin kinases with varying sequence identity from the results obtained from the BLAST search and constructed a phylogenetic tree (Figure 1A). The phylogeny of archaeal riboflavin kinases showed that the thermophilic and the mesophilic riboflavin kinases separate into distinct clades with a few exceptions in both groups. *M. maripaludis* S2 riboflavin kinase (*Mmp*RibK) was one such exception – the sequence of *Mmp*RibK and *Mj*RibK are 50.4% identical inspite of *Mmp*RibK belonging to a mesophilic archaea (Figure 1A, Table S1). Based on these results, we selected *Mmp*RibK as our candidate mesophilic CTP-dependent riboflavin kinase for further analysis.

**Figure 1:**
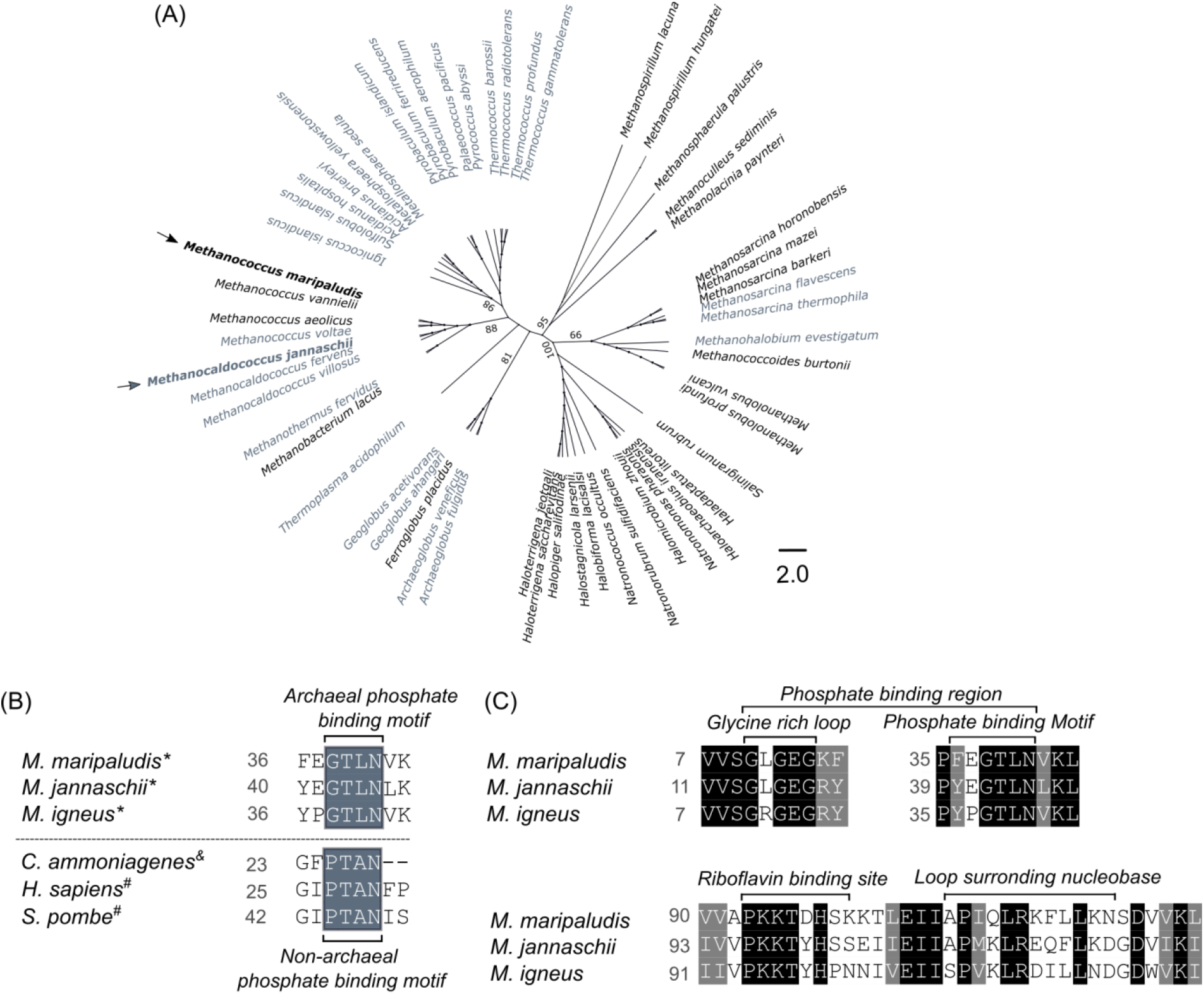
Phylogeny and multiple sequence alignment of riboflavin kinases. **(A)** We constructed a phylogenetic tree of archaeal riboflavin kinase homologs. The grey entries are riboflavin kinase sequences from thermophilic/hyperthermophilic organisms and black entries are from mesophilic organisms. The previously characterized thermophilic *Methanocaldococcus janaschii* RibK (*Mj*RibK) and the mesophilic *Methanococcus maripaludis* RibK (*Mmp*RibK) which we characterize in this study are highlighted with arrows. **(B)** Multiple sequence alignment of archaeal (indicated by *) and non-archaeal riboflavin kinases (^&^bacteria, ^#^eukaryotoes) showing the phosphate binding consensus motif present near the N-terminus– GTLN in the case of archaea and PTAN in the case of bacteria and eukaryotes. **(C)** The phosphate-binding domain, riboflavin binding site, and the region surrounding the nucleobase show well conserved patterns across archaeal sequences.

NCBI conserved domain search shows that *Mmp*RibK belongs to the conserved protein domain family pfam01982/ PRK14132, currently annotated as CTP-dependent riboflavin kinase and involved in cofactor transport and metabolism. Interestingly, some homologs in this superfamily have DNA-binding domains and they act as bifunctional transcriptional regulators/ riboflavin kinases.^[5]^

Comparing sequences of archaeal riboflavin kinase with their bacterial and eukaryotic homologs shows that these are two completely distinct types of enzymes, differing even in the conserved motif responsible for triphosphate binding (GTLN in archaeal versus PTAN in non-archaeal riboflavin kinases, as also noted previously (Figure 1B, Figure S1A).^[7,11]^ However, among the archaeal riboflavin kinases, a large degree of conservation is observed especially within the catalytic domains, as evidenced by the sequence alignment of *MmpRibK* with other homologous archaeal enzymes (Figure 1C, Figure S1B). The archaeal riboflavin kinases contain the GTLN domain and the glycine-rich GxGEG loop, responsible for nucleotide recognition and binding, mainly via the phosphate group. Another interesting feature is that the conserved amino acid sequence LRxxxL forms a loop with a small α-helix around the nucleobase portion of CTP which is not present in the corresponding loop of ATP-dependent riboflavin kinases. The nucleobase cytosine makes backbone interactions with Leu and Arg residues of the LRxxxxL domain. Finally, the conserved riboflavin binding domain PxxTxx is found to have mainly backbone interactions with riboflavin.^[7]^

### The *Mmp*RibK riboflavin kinase is functional under mesophilic conditions

We cloned, expressed and purified the enzymes *Mj*RibK and *Mmp*RibK (Figure S2 A, B). To test whether *Mmp*RibK is a functional enzyme, we conducted an assay with conditions similar to what was previously reported for *Mj*RibK, except that the temperature was set at 37°C.^[2]^ No conversion of riboflavin was observed. However, when 10 mM MgCl_2_ was added, we found that the *Mmp*RibK enzyme successfully catalyzes the conversion of riboflavin into FMN using CTP as phosphate donor, as indicated by fluorescence-TLC and verified by HPLC analysis (Figure 2A, B).

**Figure 2:**
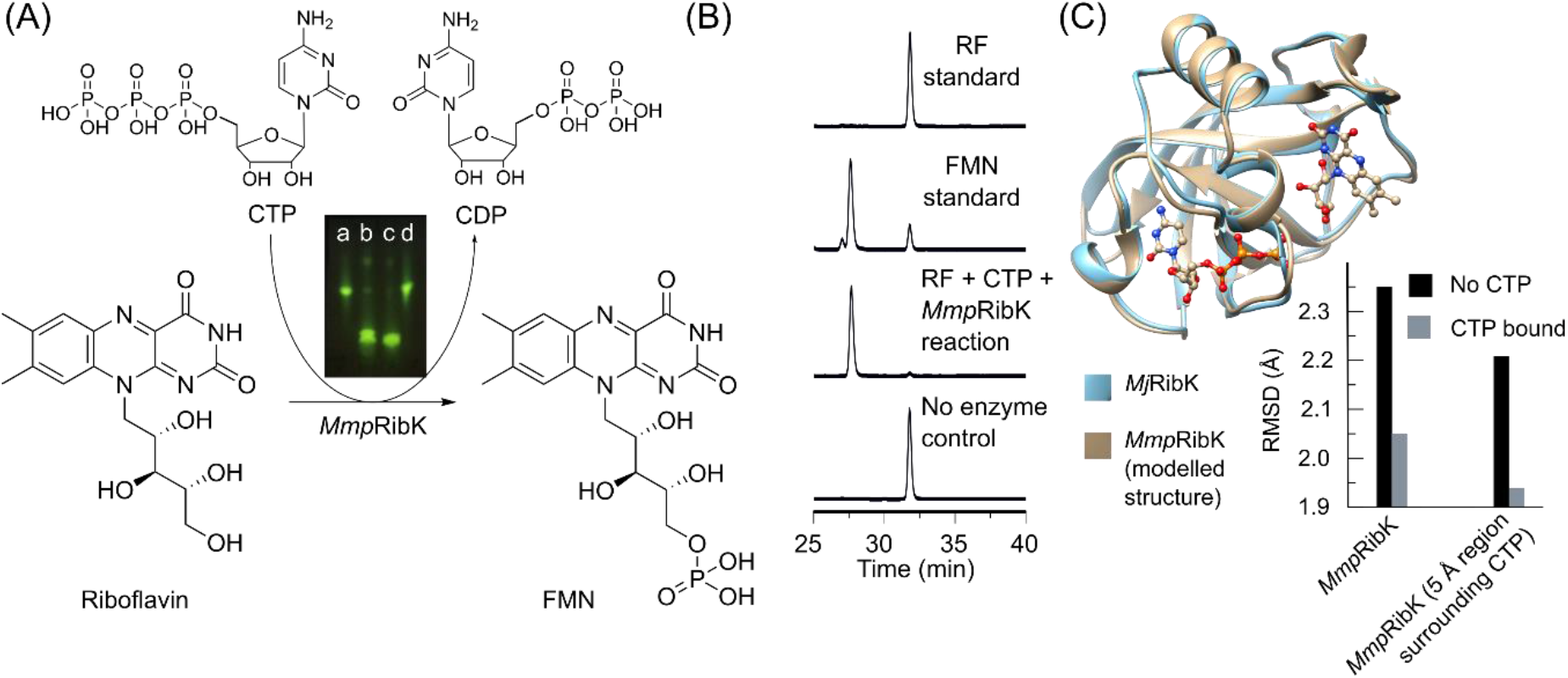
Reconstitution of the riboflavin kinase activity of *Mmp*RibK, modelling the *Mmp*RibK structure, and substrate docking. **(A)** Reaction catalyzed my *Mmp*RibK to convert riboflavin into FMN. TLC shows the fluorescent spots for (a) riboflavin standard, (b) FMN standard, (c) full reaction, and (d) No enzyme control (d). **(B)** HPLC chromatogram for the formation of FMN by *Mmp*RibK. **(C)** Overlay of the *Mj*RibK crystal structure (blue) and the modelled *Mmp*RibK structure (tan) reveals a high degree of tertiary structure similarity. *Mmp*RibK is further docked with substrates riboflavin and CTP. The inset shows the RMSD calculated for */mp*RibK in the presence and absence of the ligand CTP. The binding of the CTP appears to stabilize the enzyme.

Next, we compared some basic features of the *Mmp*RibK and *Mj*RibK homologs. First, we carried out reactions with both homologs using different nucleotide triphosphates. We found that *Mmp*RibK shows very high specificity for utilizing CTP and does not form FMN with other nucleotides such as ATP, GTP and UTP. On the other hand, *Mj*RibK shows the highest specificity with CTP, but shows 10% and 6% conversion of riboflavin to FMN with UTP and ATP, respectively (Figure S3 A, B). Temperature-wise, when held at 100°C for 5 min and then used in an assay, *Mmp*RibK is completely inactivated and shows no activity while *Mj*RibK has no significant change in activity. Lastly, as compared to *MjRibK* which is unique in its ability to catalyze the phosphate transfer reaction in absence of any divalent metal ion, *Mmp*RibK requires Mg^2+^ ion for its catalytic activity (Figure S4A).

### The modeled *Mmp*RibK structure shows high similarity with the *Mj*RibK crystal structure

The sequence of *Mmp*RibK was subjected to homology modeling, and the modeled apo-*Mmp*RibK was established to have high similarity to the crystal structure *Mj*RibK (PDB: 2VBV) (see supplementary information section for modeling details). Next, we docked the ligands CTP and riboflavin into the *Mmp*RibK structure and conducted energy minimization to obtain the most favourable configuration. Figure 2C shows the overlap of the docked *Mmp*RibK with *Mj*RibK containing CTP and riboflavin (see supplementary information section for docking details). Both the apo- and docked *Mmp*RibK models show >90% of residues are in the most favoured regions of the Ramachandran plot, which indicates that these are high quality models (Figure S5 A, B, Table S2).^[12]^ The overall quality factor of the modeled structures, calculated on the basis of statistics of non-bonded atomic interactions and distribution of atoms using ERRAT, were found to be 68.9% and 82.4% for the apo- and docked models of *Mmp*RibK, respectively, as compared to 100% for the crystal structure of *Mj*RibK. (Figure S5 C,D, E).^[13]^ Interestingly, 1 μs long molecular dynamics (MD) simulations of the whole protein *Mmp*RibK as well as that of the CTP-binding site show that the root mean square deviation (RMSD) decreases sharply when CTP and riboflavin are bound in the model, as compared to the apo *Mmp*RibK structure (Figure 2C inset). Theoretically, this indicates that *Mmp*RibK adapts a compact conformation in the presence of CTP as compared to when it is not present. Our findings were verified experimentally by a thermal shift assay, in which the broad signal of the apo wild-type *Mmp*RibK changed into a sharp depression on adding CTP (Figure S2C).^[14]^ The control reaction, where ATP was added, showed no change. This not only aligns with our previous observation that the enzyme *Mmp*RibK does not catalyse the riboflavin kinase reaction with ATP, but also indicates that ATP is likely unable bind in the enzyme’s nucleotide binding site.

### *Mmp*RibK demonstrates greater molecular flexibility and lower stability under a range of temperatures as compared to *Mj*RibK

We began our functional analysis of the enzyme *Mmp*RibK by testing its activity at varying temperatures. *Mmp*RibK was found to be most active in the temperature range of 40°C-50°C, after which its activity declined until it was completely inactivated beyond 70°C (Figure 3A). In a similar experiment with *Mj*RibK, we observed the optimal activity extending from 45°C to 100°C (Figure 3A). Next, the stability of *Mmp*RibK and *Mj*RibK enzymes were probed using a chemical denaturation agent guanidium hydrochloride (Gdn-HCl).^[15]^ Both enzymes were subject to varying concentrations of Gdn-HCl and their activities were analyzed using fluorescence-TLC. We observed that *Mmp*RibK tolerates concentrations of Gdn-HCl only up to 1.2 M, while there is no significant change is activity of *Mj*RibK up to 2.5 M Gdn-HCl (Figure S6 A, B). Both these experiments indicate that even though *Mmp*RibK and *Mj*RibK sequences share 50.4% identity, the differences in their amino acid compositions and consequently structures confer a large degree of rigidity to *Mj*RibK. To gain a better understanding of the molecular flexibility of both enzymes, we conducted MD simulation studies on the substrate-bound *Mj*RibK and *Mmp*RibK structures and calculated the root mean square fluctuation (RMSF) of the enzyme at 27°C and 72°C (see simulation details in the supplementary information section). The RMSF for *Mmp*RibK at 72°C increases significantly as compared to that at 27°C, indicating that temperature increase makes *Mmp*RibK more flexible, consequently resulting in destabilization of the bound CTP (Figure 3B). In sharp contrast, we find that the RMSF of *Mj*RibK remains similar at temperatures of 27°C and 72°C. Taken together, our experimental results and MD simulations strongly suggest that there exist features within the *Mj*RibK enzyme which confer it with the ability to retain its function over a large range of temperatures as compared to *Mmp*RibK.

**Figure 3:**
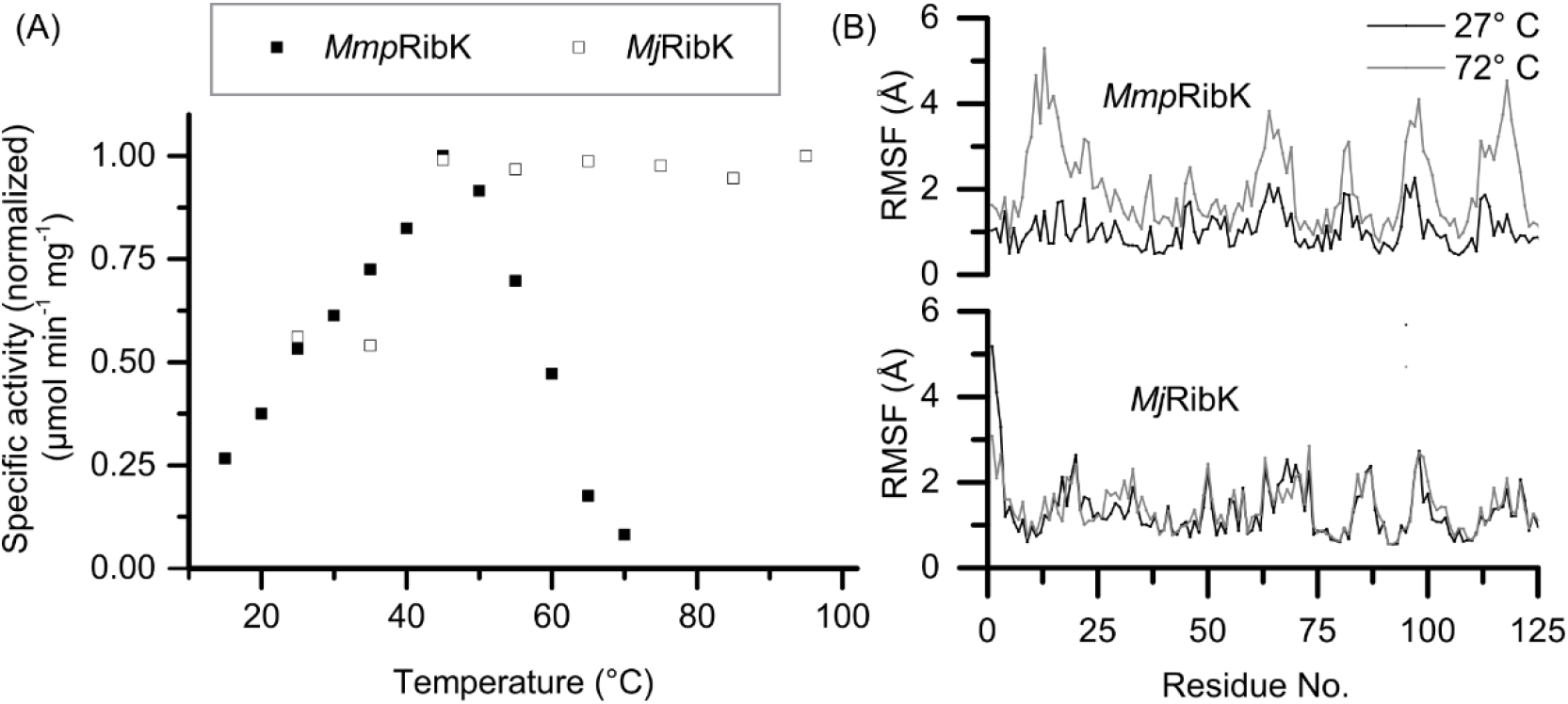
Temperature dependence of *Mmp*RibK and *Mj*RibK: **(A)** The specific activity of *Mmp*RibK and *Mj*RibK was calculated from 15°C to 70°C and the normalized data is plotted as a function of temperature. The activity increases for *Mmp*RibK and peaks at 45°C after which it decreases gradually until the enzyme is no longer active at 70°C. In contrast, *Mmp*RibK shows activity at temperatures ranging from 45°C to 95°C. **(B)** Root mean square fluctuation (RMSF) of *Mmp*RibK and *Mj*RibK were computationally evaluated at 27°C and 72°C. *Mmp*RibK shows elevated RMSF at 72°C indicating that it is more flexible as compared to *Mj*RibK, which is stable at both temperatures.

### Establishing optimal metal dependence, pH range, and salt concentration for *Mmp*RibK activity

We have already established that *Mmp*RibK, unlike *Mj*RibK, requires a metal ion to function as a kinase.^[2]^ *Mmp*RibK showed no activity in the absence of a metal, however, it showed full turnover when the divalent metal ion Mg^2+^ was added into the reaction (Figure S4A). We further examined the range of divalent metal ions Mg^2+^, Zn^2+^, Mn^2+^, Ni^2+^, Co^2+^, Fe^2+^, Cu^2+^, and Ca^2+^, and found that *Mmp*RibK is able to phosphorylate riboflavin most effectively in the presence of Mg and Zn (Figure S4B). The overall trend of activity with different divalent metal ion is Mg^2+^ = Zn^2+^>Mn^2+^>Ni^2+^>Co^2+^>Fe^2+^>Cu^2+^> Ca^2+^ (Figure S4B). This preference is reminiscent of the rat liver riboflavin kinase which is reported to exhibit a preference of Zn^2+^ ion followed by Mg^2+^ ion.^[16]^

Next, we tested the *Mmp*RibK enzyme reaction in the pH range of 4.3-12 to determine the optimal pH for its activity. We observed a steady increase in the activity of the enzyme up to pH 11 (Figure S4C). A similar trend where increase in pH leads to higher activity has been previously noted for the *Neurospora crassa* riboflavin kinase.^[17]^

While conducting enzyme assays with *Mmp*RibK and *Mj*RibK, we observed that the activity of *Mmp*RibK was slower for the conversion of riboflavin into FMN in comparison to *Mj*RibK. As *M. maripaudis* was first isolated from a salt marsh environment, and such ecosystems contain high concentrations of NaCl, we hypothesized that the *Mmp*RibK activity might be improved in the presence of NaCl. Indeed, the enzymatic activity improved with added NaCl, and a maximum of 2.3x-fold increase was observed with 2M NaCl (Figure S4D). To our knowledge, this is the first riboflavin kinase which has been shown to require a high concentration of NaCl.

### Steady-state kinetic analysis of *Mmp*RibK

We conducted the steady-state kinetic analysis of *Mmp*RibK for both substrates riboflavin and CTP in the presence of NaCl. In the presence of saturating concentration of CTP, riboflavin displays the follows kinetic parameters: V_max_ 5.9 μM min^−1^, K_m_ 185.8 μM and *k_cat_* 0.19 s^−1^ (Figure 4A, Figure S7 A). Under saturating concentration of riboflavin, the kinetic parameters for CTP are: V_max_ 3.9 μM min^−1^, K_m_ 236.0 μM and *k*_cat_ 0.14 s^−1^ (Figure 4B, Figure S7 B). We compared these values to the parameters reported for *Mj*RibK and established that overall, both enzymes demonstrate similar reaction kinetics, even though there are small differences in reaction conditions for optimal activity (Table 1).^[2]^

**Figure 4:**
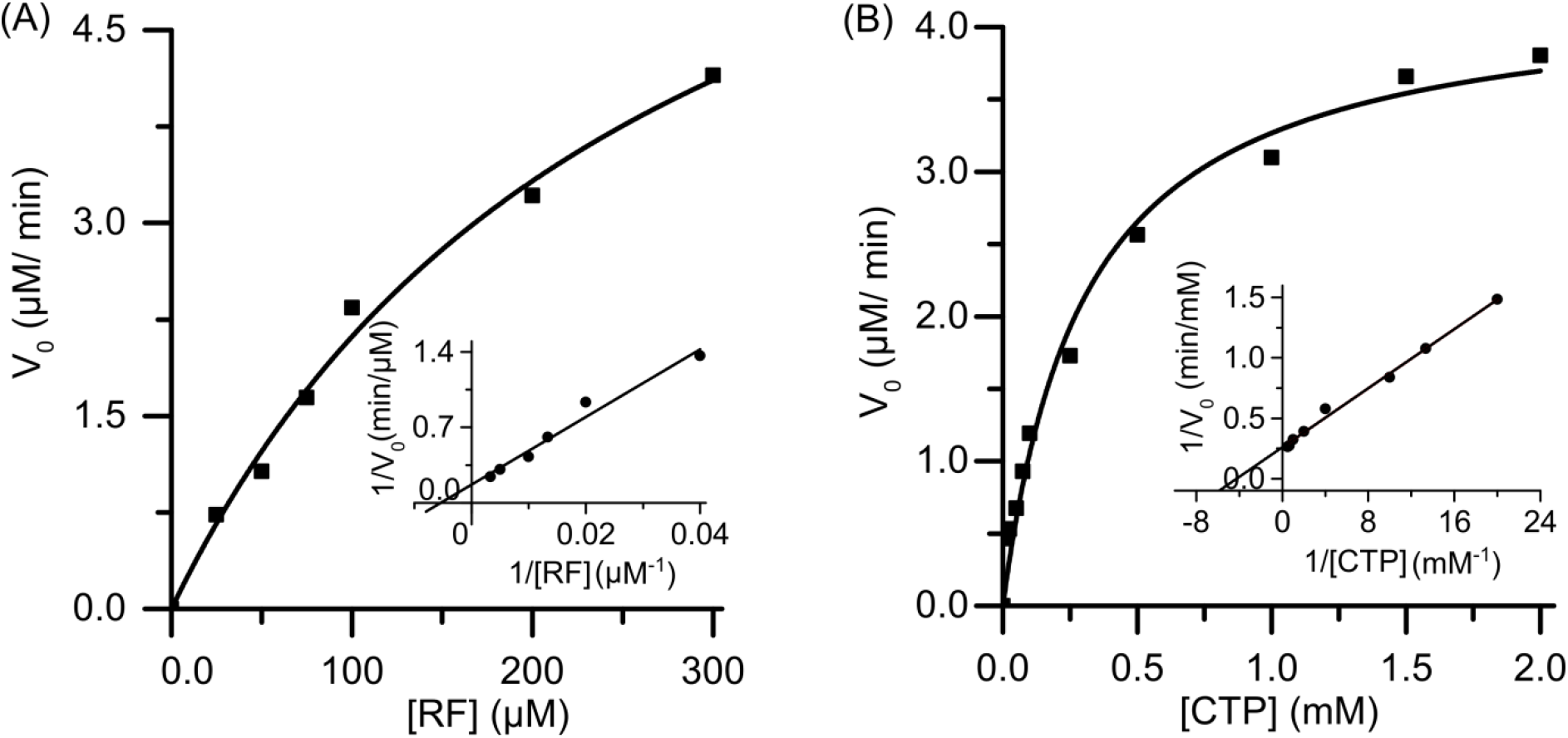
Steady-state kinetics of *Mmp*RibK with substrates riboflavin and CTP: **(A)** Michaelis-Menten plot of *Mmp*RibK for riboflavin under saturating conditions of CTP. The same data is represented as a Lineweaver-Burk plot, shown in the inset. **(B)** Michaelis-Menten kinetics of *Mmp*RibK for CTP under saturating conditions of riboflavin. The Lineweaver-Burk plot for the same data set is provided in the inset.

**Table 1:** Kinetics parameters for *Mmp*RibK and *Mj*RibK

|  | *MmpRibK* (this paper) |  | *MjRibK*<sup>2</sup> (values obtained from Mashhadi et al., JBact, 2008) |  |
| --- | --- | --- | --- | --- |
|  | Riboflavin | CTP | Riboflavin | CTP |
| $V_{\max}$ ($\mu\text{M min}^{-1}$) | 5.9 | 3.9 | Data not available | Data not available |
| $K_m$ ($\mu\text{M}$) | 185.8 | 236.0 | 159 | 180 |
| $k_{\text{cat}}$ ($\text{s}^{-1}$) | 0.19 | 0.14 | 0.334 | 0.342 |

In summary, we have established that *Mmp*RibK is a functional mesophilic riboflavin kinase with a strong preference for CTP over other nucleotide triphosphates, with temperature, pH and NaCl concentration optima of 45°C, 11 and 2 M, respectively.

### Analysis of the molecular basis of thermostability of *Mj*RibK and rational engineering of the *Mmp*RibK sequence to introduce thermostable elements

Our results have established that *Mmp*RibK differentiates itself adequately from its thermophilic homolog *Mj*RibK while sharing a high degree of sequence identity and structural features. Thus, this pair of CTP-utilizing kinase enzymes is an excellent candidate for analysing the general principles of thermal stability in enzymes via rational engineering.

We conducted a computational analysis as described in the methods section to predict salt bridges in the *MmpRibK* and *MjRibK* structures. The *Mj*RibK had more salt bridge interactions than *Mmp*RibK, including one within the nucleobase loop region in its active site (Table 2, Figure 5 A-C). This is expected, as salt bridges are noted to confer rigidity and subsequently thermostability to proteins.^[18]^ In order to test the contributions of these predicted salt bridges, we introduced the residues that formed additional salt bridge in *Mj*RibK into the corresponding positions within the *Mmp*RibK sequence, and created three mutants, Mut 1-3 (Table3, Figure 5 A-C). Additionally, proline residues are also known to confer thermostability in a mesophilic enzyme by rigidifying the overall structure.^[19]^ Based on the sequence alignment of *Mj*RibK and *Mmp*RibK, we replaced residues T21 and H22 in *Mmp*RibK by two consecutive prolines to create Mut4 (Table3, Figure 5D). Mut 1-4 were tested for activity at 60 °C, where the wild-type enzyme drastically loses activity. Except Mut 1, the remaining three mutants showed similar or better turnover as compared to the wild-type enzyme. Hence, we proceeded to create combinations of Mut 2-4 involving salt bridges and proline residues to generate Mut5-8. (Table 3).

**Figure 5:**
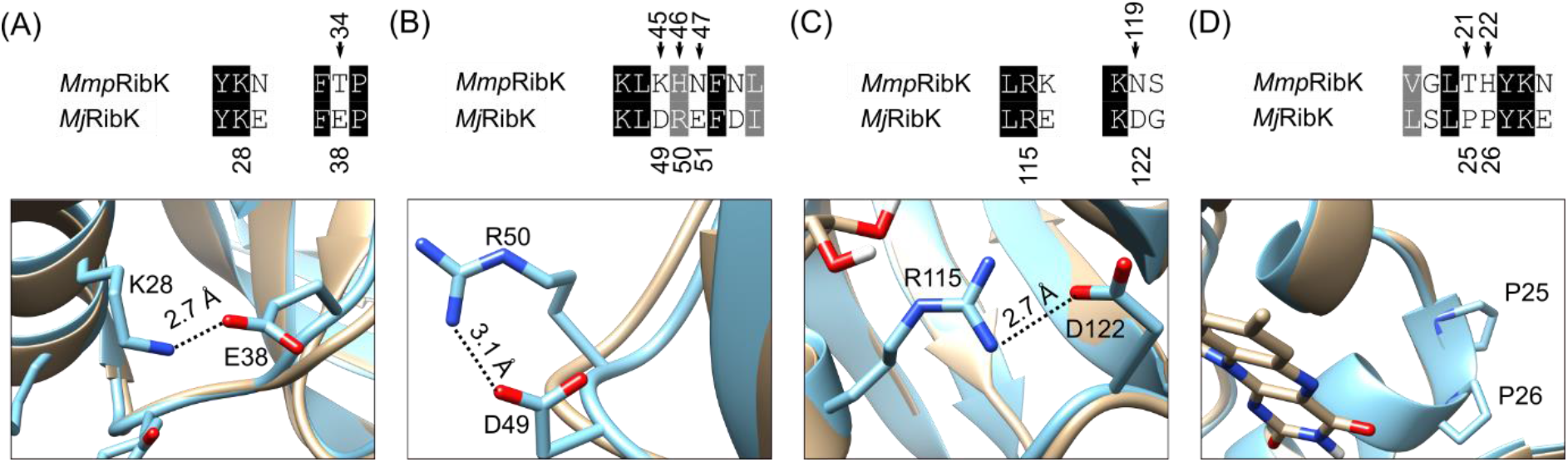
Overlay of modeled structure of *Mmp*RibK (tan) and crystal structure of *Mj*RibK (blue) depicting the salt bridges and non-conserved proline residues considered for mutation. **(A)** K28-E38 salt bridge in *Mj*RibK corresponding to *Mmp*RibK Mut1 (T34E) **(B)** D49-R50 salt bridge in *MjRibK* corresponding to *Mmp*RibK Mut2 (KHN45-47DRE) **(C)** R115-D122 salt bridge corresponding to *Mmp*RibK Mut3 (N119D) **(D)** Non-conserved proline residues in *Mj*RFK corresponding to Mut4 (TH21-22PP)

**Table 2:** Salt bridges present in *MjRibK* (data obtained from the crystal structure PDB: 2VBU with ligands CDP and Mg^2+^).

| No. | Salt bridges in *MjRibK* | Corresponding residues in *MmpRibK* |
| --- | --- | --- |
| 1 | **E38-K28\*** | T34-K24 |
| 2 | **E38-K32\*** | T34-E28 |
| 3 | E41-K28 | E37-K24 |
| 4 | **D49-R50\*** | K45-H46 |
| 5 | E61-K97 | E57-K93 |
| 6 | **D90-K81\*** | N87-K81 |
| 7 | **D122-R115\*** | N119-R111 |
| 8 | D124-K121 | D120-K117 |
| *Salt bridges that exist in *MjRibK* but are absent in *MmpRibK* |  |  |

**Table 3:** List of mutants and corresponding mutations

| Mutant | Residue/s of *MmpRibK* replaced with that in *MjRibK* | Putative salt bridge introduced | No. of salt bridges introduced | No. of residues replaced by Pro |
| --- | --- | --- | --- | --- |
| Mut1 | T34E | E34-K24 | 1 | 0 |
| Mut2 | KHN(45-47)DRE | D45-R46 | 1 | 0 |
| Mut3 | N119D | D119-R111 | 1 | 0 |
| Mut4 | TH(21-22)PP | - | 0 | 2 |
| Mut5 | KHN(45-47)DRE/N119D | D45-R46/<br>D119-R111 | 2 | 0 |
| Mut6 | KHN(45-47)DRE/TH(21-22)PP | D45-R46 | 1 | 2 |
| Mut7 | TH(21-22)PP/N119D | D119-R111 | 1 | 2 |
| Mut8 | KHN(45-47)DRE/N119D/TH(21-22)PP | D45-R46/<br>D119-R111 | 2 | 2 |

### Molecular dynamics simulations, thermal shift analysis and specific activity of the *Mmp*RibK mutants validate the role of amino acids that confer thermostability

To gain a better understanding of the feasibility and stability of the eight *Mmp*RibK mutants, we conducted MD simulation studies and calculated the RMSD of their modeled structures over a period of 300 ns, followed by the RMSF of individual residues in each mutant (see simulation details in the supplementary information section) (Figure 6 A, B). We found that all eight enzymes exhibited a relatively stable RMSD value except for Mut 4 and Mut 8 (Figure 6A). Also, the RMSF values at 72 °C indicate that the residues in all the mutants remain stable except those in Mut 1. This behavior of Mut1 compares to the wild-type enzyme at that temperature, confirming our experimental result that Mut1 loses activity at 60 °C or higher (Figure 6B, C). We performed a thermal shift analysis with wild-type *Mmp*RibK and its mutants in order to establish their melting temperatures. Briefly, the change in the signal intensity with increasing temperature, attributable to the gradual unfolding of the protein mixed with a fluorescent dye, was correlated to its thermal stability (Figure 6C). Our results show that Mut1 (with an E-K salt bridge) has a similar melting temperature as the wild-type enzyme, Mut4 (with two P residues) was stabilized to a small extent (T_m_= 38.8°C) and Mut2 (with a D-R salt bridge) and Mut3 (with a D-R salt bridge) were significantly stabilized (T_m_=46.4°C and T_m_=47°C, respectively) (Figure 6C, Figure S8, Table 3). The remaining combination mutants Mut 5-8 all displayed T_m_ higher than the wild-type enzyme, but surprisingly, these were lower than that of Mut 2 and Mut 3 (Figure 6C, Figure S8).

**Figure 6:**
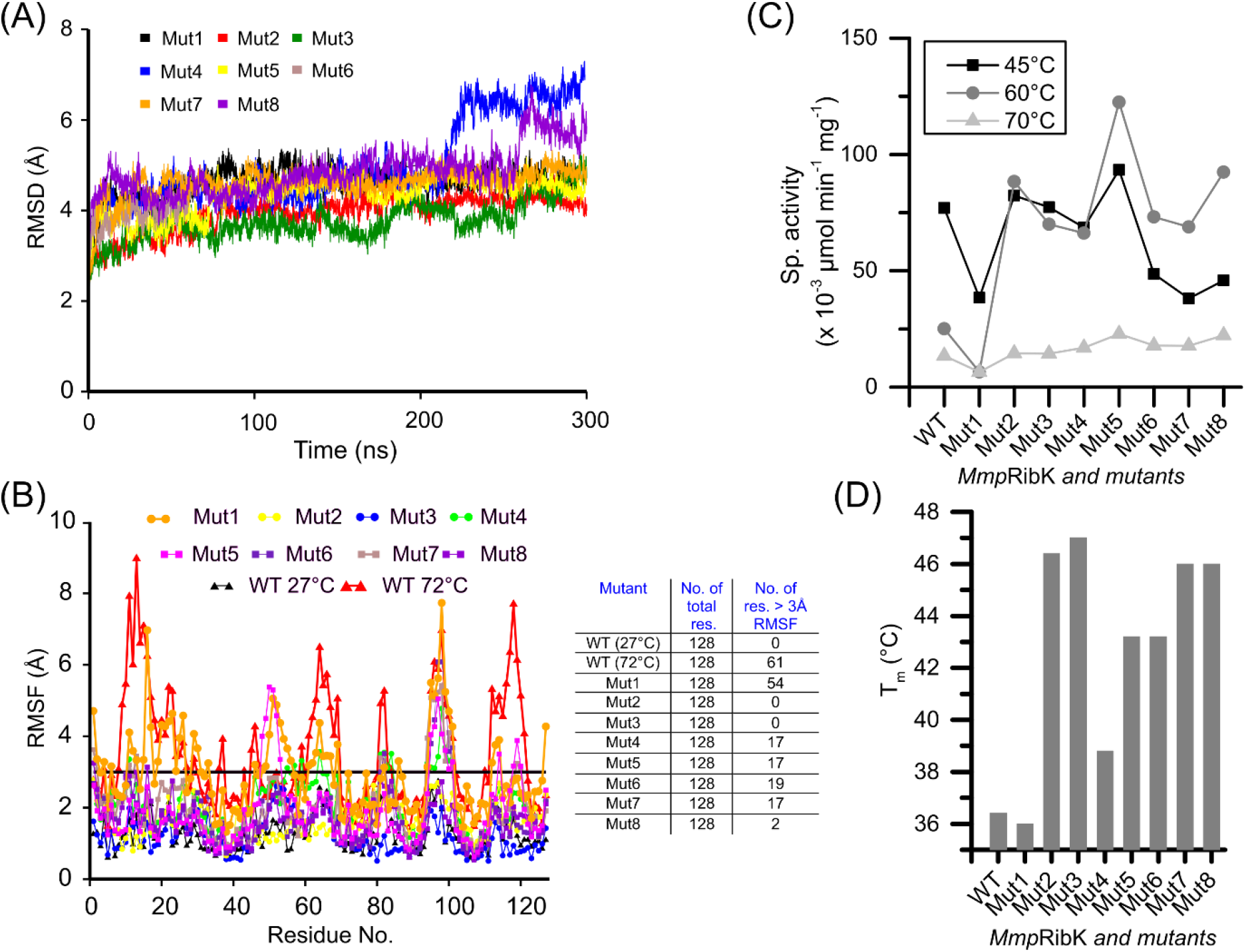
RMSD and RMSF calculations, specific activity and melting temperature analysis of the wild-type and *Mmp*RibK mutants. **(A)** RMSD of the wild-type enzyme and the eight MmpRibK mutants was calculated over a 300 ns long molecular dynamic trajectory. The RMSD analysis shows that enzymes are stable except for Mut 4 and Mut 8 that show higher values indicating that these mutants may be partially unfolded (B) An RMSF performed for the amino acid residues of the *Mmp*RibK wild-type and mutants calculated over a 300 ns long molecular dynamic trajectory, show that Mut 2-8 have gained thermostability, except Mut 1 which has high fluctuations and behaves similarly to the wild-type *Mmp*RibK at 72 °C. The table show the number of amino acid with greater than 3 Å RMSF. **(C)** The specific activities of *Mmp*RibK and mutants measured at 45°C, 60°C, and 70°C show that all the mutants except Mut1 have higher specific activity as compared to the wild-type at 60 °C **(D)** Melting temperatures of the wild-type *Mmp*RibK and its mutants determined by the thermal shift assay show higher stability in Mut2-8

Since the thermostabilization of an enzyme may occur at the cost of its activity, we compared the activity of the wild-type and the mutants at higher temperatures. Temperature-dependent activities of the wild-type *Mmp*RibK and its mutants were conducted at 45°C, 60° and 70°C (Figure 6D). Wild-type *Mmp*RibK has reduced activity at 60°C and 70°C in comparison to that at 45°C (also seen in Figure 3A). At 45°C, the activities of wild-type *Mmp*RibK, Mut2, Mut3, Mut4 and Mut5 are comparable, while Mut6, Mut7 and Mut8 have reduced activity at this temperature. On the other hand, at 60°C, all mutants except Mut1, have higher activity when compared to wild-type *Mmp*RibK. The highest activity is displayed by Mut5 which contains two added salt bridges (Table 3). Finally, at 70°C, the wild-type and mutants have significantly diminished activity. This result demonstrates that using the rational engineering strategy of comparative sequence analysis followed by creating appropriate mutants, we were able to achieve a 15°C improvement in thermal stability of *Mmp*RibK without loss in activity.

### Discussion

The cofactor biosynthesis pathway enzymes in archaea have unravelled many interesting reaction mechanisms to date, and the CTP-dependent riboflavin kinase is one such enzyme. Until now, only one thermophilic homolog *Mj*RibK had been characterized. In this study, we report the reconstitution of the activity of the first mesophilic CTP-dependent riboflavin kinase homolog *Mmp*RibK, and establish its biochemical properties.

On testing for various reaction parameters, we found that *Mmp*RibK varies in several ways from *Mj*RibK. A feature of the *Mj*RibK homolog that stands out is that it does not require a metal ion for its function.^[2]^ In contrast, we found that *Mmp*RibK requires a metal ion for activity, and gives the best activity with Mg^2+^ and Zn^2+^. As most kinases use a metal ion to stabilize the nucleotide triphosphate within its active site and to catalyse the phosphate transfer reaction, we tried to understand the role of the metal ion in archaeal RibK enzymes. To do so, we conducted an MD simulation with the enzyme and substrates in the absence and presence of MgCl_2_. Our results show that *Mj*RibK is able to bind CTP and riboflavin in the absence of the magnesium ion and the enzyme has favourable hydrogen bonding interactions with both substrates, confirming that *Mj*RibK is indeed a metal-free riboflavin kinase (Figure S9 A, B).^[2]^ In the case of *Mmp*RibK, the absence of Mg^2+^ results in weak binding of CTP in the active site with few hydrogen bonds (Figure S9 C, D). This changes to a larger number of hydrogen bonds stabilizing the CTP in the active site in the presence of Mg^2+^ (Figure S9 E, F). Additionally, a constant distance is maintained between the Mg^2+^ and the phosphate residues of CTP indicating that the metal ion is important for its stabilization (Figure S9 G). Finally, the Mg^2+^ ion appears to be snugly held in the active site, hexacoordinated to *Mmp*RibK residues Gly 14, Asn 41, Glu 104, the oxygen of riboflavin, the γ-phosphate of CTP, and an active site H_2_O.

Our choice of studying *Mmp*RibK was based on its high degree of sequence identity to *Mj*RibK, also evidenced by their proximity on the phylogenetic tree and from the similar proportions of individual amino acids that constitute each of the homologs (Figure 1A, Table S3). Interestingly, the mol% of amino acid residues significant differs for the ones that are reportedly involved in salt bridges (Arg, Lys, Asp, Glu, His) and in providing rigidity to the protein (Pro) (Figure S10).^[20]^ As expected, the thermophilic *Mj*RibK has more salt bridge-forming and rigidity-conferring residues as compared to the mesophilic *Mmp*RibK, the only exception being His.^[18]^ Our experimental observation that *Mmp*RibK functions optimally at 45 °C, a temperature much lower than *Mj*RibK, led us to probe the biochemical basis for thermal stability of *Mj*RibK using bioinformatics and computational tools. The introduction of specific thermostability elements i.e. salt bridge residues and Pro to confer rigidity at the corresponding positions in the *Mmp*RibK enzyme led to the creation of mutant variants of *Mmp*RibK that show optimal activity at 60 °C i.e. 15 °C higher than the wild-type enzyme. Lastly, as *Mmp*RibK is a riboflavin kinase that is specific for its use of CTP as its phosphate donor, we investigated whether the thermostable mutants we had created had either lost activity or had lost their specificity for CTP at 60 °C. We found that with the exception of one mutant (Mut1), all the other mutants we created had similar or higher activity than the wild-type *Mmp*RibK with CTP as a substrate even at 60 °C (Figure 6D). At the same time, the reaction of mutants with NTPs other than CTP showed that their specificities were unaltered even after mutation (Figure S11).Taken together, our results establish that our method of rational engineering the amino acids of a mesophilic protein homolog by comparing its sequence and secondary structure with the corresponding thermophilic homolog, followed by systematically introducing them in the mesophilic enzyme is a reliable general strategy to produce thermostable enzymes for various applications.

### Conclusion

In this study, we report the successful reconstitution and biochemical characterization of a novel mesophilic CTP-dependent riboflavin kinase from the archaea *M. maripaludis*. The use of CTP as the substrate for a kinase is a rare occurrence, thus the characterization of *Mmp*RibK adds to the handful of examples in literature that will allow us to explore the origins of nucleotide choice in kinases. Our studies on the characterization of *Mmp*RibK and creation of its thermostable mutants not only adds important mechanistic information to this class of enzymes, but also puts forth an important candidate for biochemical and structural investigations and bioengineering applications in the future.

## MATERIALS AND METHODS

### Materials

Genomic DNA of *Methanocaldococcus janaschii* DSM 2661 and *Methanococcus maripaludis* S2 were a gift from Biswarup Mukhopadhyay at Virginia State University. The primary assay reagents riboflavin and FMN are from Sigma-Aldrich, and the nucleotide triphosphates were obtained from Jena Biosciences. All other reagents and media components used in cloning, protein expression and purification and enzyme assays were obtained from Sigma-Aldrich, TCI chemicals, Rankem, SRL chemicals and Himedia, unless otherwise stated. Polymerase chain reaction (PCR) reagents and kits were from Agilent and Himedia, and the high fidelity PrimeSTAR GXL DNA polymerase from DSS Takara Bio USA was used for cloning. Sypro-orange dye used in the thermal shift assay was obtained from Thermofisher Scientific.

### Bioinformatic and phylogenetic analysis for selection of *Mmp*RibK as the candidate

To identify a suitable mesophilic CTP-dependent riboflavin kinase homolog for our studies, the FASTA protein sequence of *Mj*RibK (Swiss-Prot accession number Q60365) was used to perform a <u>B</u>asic <u>L</u>ocal <u>A</u>lignment <u>S</u>earch <u>T</u>ool (BLAST) analysis, which yielded 1813 distinct CTP-dependent riboflavin kinase entries^[21–23]^. A phylogeny study was then conducted for a curated set of 56 protein sequences of archaeal thermophilic and mesophilic RibK enzymes from the list of BLAST search candidates. To do so, we created a multiple sequence alignment using the **MU**ltiple **S**equence **C**omparison by **L**og-**E**xpectation (MUSCLE) alignment tool, which revealed that some of the riboflavin kinase proteins contained a transcriptional regulator.^[24,25]^ The transcriptional regulator domains were removed using BioEdit, so as to only consider the riboflavin kinase domain sequence for analysis and the alignment was saved as a phylip4.0 file. The alignment was then subjected to maximum likelihood/ rapid bootstrapping analysis on RaXML-HPC2 on XSEDE on the CIPRES gateway to produce a phylogenetic tree, which was visualized using Figtree.^[26]^Two additional multiple sequence alignments of only archaeal riboflavin kinases and archaeal and bacterial riboflavin kinases were also conducted using the method described above.

### Molecular cloning, overexpression and purification

*MjribK* and *MmpribK*, the riboflavin kinase genes from *Methanocaldococcus jannaschhi* and *Methanococcus maripaludis* S2, respectively, were amplified using PCR and inserted between the NdeI and BamHI restriction sites of pET21a plasmid vector using restriction-free cloning.^[27]^ The plasmids containing the *MjribK* and *MmpribK* genes were transformed into *E. coli* BL21(DE3) competent cells and an expression check was conducted to verify whether the corresponding proteins were being expressed. Autoclaved Luria Bertani broth supplementated with 50 μg/mL of kanamycin was inoculated with 1% of overnight grown primary culture and left at 37°C inside the incubator shaker. When the OD600 of culture reached 0.6-0.8, protein expression was induced with 1 mM isopropyl β-D-1-thiogalactopyranoside (IPTG) at 18°C and left to shake for 15 h, after which the cells were centrifuged to pellets and stored at −80 degrees. Purification of enzyme having N-terminal hexahistidine tag was done by Ni-NTA affinity chromatography using an Akta pure FPLC system (GE Healthcare). Equilibration buffer was 50 mM tris-HCl pH 8.0, 300 mM NaCl, 20 mM imidazole and 0.025% β-mercaptoethanol. Protein was eluted with buffer containing 50 mM Tris-HCl pH 8.0, 300 mM NaCl, 20 mM imidazole 250 mM imidazole and 0.025% β-mercaptoethanol. The concentration of protein was checked using Bradford assay using BSA as standard.^[28,29]^ All mutants of *Mmp*RibK were cloned, expressed and purified in the same manner. The primer sequences used for the cloning of wild-type and mutants and the plasmids generated are listed in the supplementary information Table S4 and S5, respectively. The purity and molecular weight of the enzymes were analyzed by 15% SDS-PAGE gel electrophoresis (Figure S2 A). The wild-type *Mmp*RibK enzyme was also subjected to Matrix-Assisted Laser Desorption/ Ionization – Time of Flight (MALDI-TOF) analysis as per reported protocols and its mass was verified to be 16.37126 kDa (corresponding to the calculated mass of the *Mmp*RibK protein with the hexahistidine tag) (Figure S2B).

### Enzyme assay and chromatographic analysis

We tested the enzymatic activity of wild-type *Mmp*RibK using 300 μM riboflavin, 4 mM CTP, 10 mM MgCl_2_, 100 mM Tris-HCl pH 8.0 and 1 μM enzyme, modifying on previous reported conditions for the assay of *Mj*RibK.^[2]^ The reaction mixture was incubated at 37°C overnight and visualized using fluoresce-based thin layer chromatography (TLC) (Merck Silica gel 60 F254) using the long wavelength (365 nm) of a handheld UV lamp (Spectroline E-series) with the solvent system water: acetic acid: n-butanol in the ratio 5:1:4.^[30]^ The retention time of riboflavin and FMN in this solvent system was 0.45 and 0.2, respectively. The enzymatic reaction product was further analyzed by high performance liquid chromatography (Agilent HPLC system (1260 Infinity II) equipped with UV-Vis DAD detector on a C-18 reverse-phase column (Phenomenex, Gemini 5μm NX-C18 110 Å, 250 × 4.6 mm). The HPLC method used for analysis was derived from the one previously used to assay *Mj*RibK.^[2]^ The mobile phases used were 10 mM ammonium acetate pH 6.5 (Solvent-A) and methanol (Solvent-B). Gradient elution of analytes were done at 0.5 mL/min as follows. 0-5 min 95% A, 5-40 min from 95% A to 20% A, 40-60 min from 20% A to 95% A. The analytes were detected at 260 nm, 280 nm, 365 nm and 440 nm. The mutants of *Mmp*RibK were assayed and analyzed in a manner identical to the wild-type enzyme, unless otherwise mentioned.

The functional characterization and optimization of *Mmp*RibK using NaCl, alternate nucleotides, different metal ions, varying temperatures pH and guanidinium hydrochloride assisted chemical denaturation were conducted as described in the supplementary information.

### Determination of kinetic parameters for *Mmp*RibK for the substrates riboflavin and CTP

The kinetic parameters for riboflavin - K_m_, *k_cat_*, V_max_ - were determined using 4 mM CTP. The concentration of riboflavin was varied from 25 μM to 300 μM. Similarly, to determine the kinetic parameters for CTP, 300 μM of riboflavin was used and CTP concentration has been varied from 12.5 μM to 2 mM. All kinetic experiments have been done in the presence of 2M NaCl. The concentration of the product FMN synthesized was determined at various time points from the area under the curve obtained from the HPLC chromatogram using a previously generated standard curve of FMN. From the plot of the rate, initial velocities at different concentrations are determined. The Michaelis-Menten and Lineweaver-Burk data were plotted with Origin 9, using nonlinear fitting with the Hills algorithm, 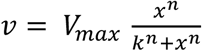 where n is the number of cooperative sites. For single-substrate model, n is fixed to 1.

### Structural analysis and ligand docking

The PDB structure of the wild-type and mutant *Mmp*RibK enzymes were modelled using Modeller.^[31]^ The stereochemical quality of the modelled structures was estimated using programs PROCHECK and ERRAT.^[12,13]^ All structural analysis were done using molecular visualization softwares ‘Visual Molecular Dynamics’ (VMD) and UCSF Chimera.^[32,33]^ The residues involved in the formation of salt bridges were identified using the salt bridges analysis tool of VMD. Docking of the ligands riboflavin and CTP are done using Autodock 4.^[34,35]^

### Molecular Dynamics simulations

We docked CTP and riboflavin in the *Mmp*RibK structure and then performed a molecular dynamics (MD) simulation. We used AMBER99SB force field for the protein.^[36]^ To create the amber force field of CTP and riboflavin we used antechamber module of AmberTool.^[37]^ First, we conducted a quantum mechanical optimization of the CTP using HF/6-31G* basis set in GAUSSIAN03 software following which restricted electrostatic potential (RESP) charges on the atoms of CTP and riboflavin were calculated.^[38]^ The topology and coordinates generated from AmberTools were converted into the GROMACS format using a perl program amb2gmx.pl. We put the CTP-riboflavin bound protein in a cubic box of length 70 Å. Then, the system was solvated with ~2000 TIP3P water molecules. Further, we added 150 mM MgCl_2_ ion solution after which we optimized the system using steepest descent method, followed by heating the system to 27°C using the Berendsen thermostat with coupling constant of 2 ps (NVT condition).^[39,40]^ After that, we have performed 4 ns NPT simulation to adjust the box size. Finally, we performed production run for 300 ns at 27°C and at 72°C using Nose-Hoover thermostat.^[41]^ All the analysis is done over production run. To study the effect of MgCl_2_ we performed two set of simulations of *Mmp*RibK-CTP-riboflavin complex in (i) absence and (ii) presence of Mg^2+^. All the simulations were performed using GROMACS simulation package (See supplementary material for details).^[42]^

### Thermal shift assay

The thermostability of mutants has been tested using thermal shift assay.^[14]^ Sypro-orange dye (5000x commercial stock) was added at final concentration of 2.5x to 5 μM of the wild-type *Mmp*RibK and its mutants (total volume of 50 μL in individual wells of a 96-well plate). The change in the fluorescence intensity during gradual unfolding owing to rise in the temperature is measured using CFX Real Time PCR instrument (Bio-Rad). The negative derivative of the change in fluorescence intensity with temperature was plotted against temperatures, of which the lowest value corresponds to the T_m_ of enzyme.

## Acknowledgements

We are grateful to Dr. Biswarup Mukhopadhyay for his gift of the genomic DNA of *M. janaschii* and *M. maripaludis* used in this study. We thank Yamini Mathur, Rupali Sathe, Ateek Shah, Bappa Ghosh and Nishant Singh for their critical inputs on experiments and their execution. We thank Harinath Chakrapani, Santosh Kumar Jha, Arnab Mukharjee, and Gayathri Pananghat for insightful discussions. We thank Mohan Reddy at IISER Pune for his help with MALDI-TOF analysis and the help received from the LC-MS and MALDI-TOF facilities and their managers at IISER Pune. We also acknowledge contributions by summer student Lavisha Parab and semester project student Saswata Nayak. Y.K. acknowledges IISER Pune and University Grant Commission, India, and R.S acknowledges CSIR and the IIT Bombay postdoctoral fellowship for funding.

