## Supplementary information for "Characterization of a novel mesophilic CTP-dependent riboflavin kinase and rational engineering to create its thermostable homologs"

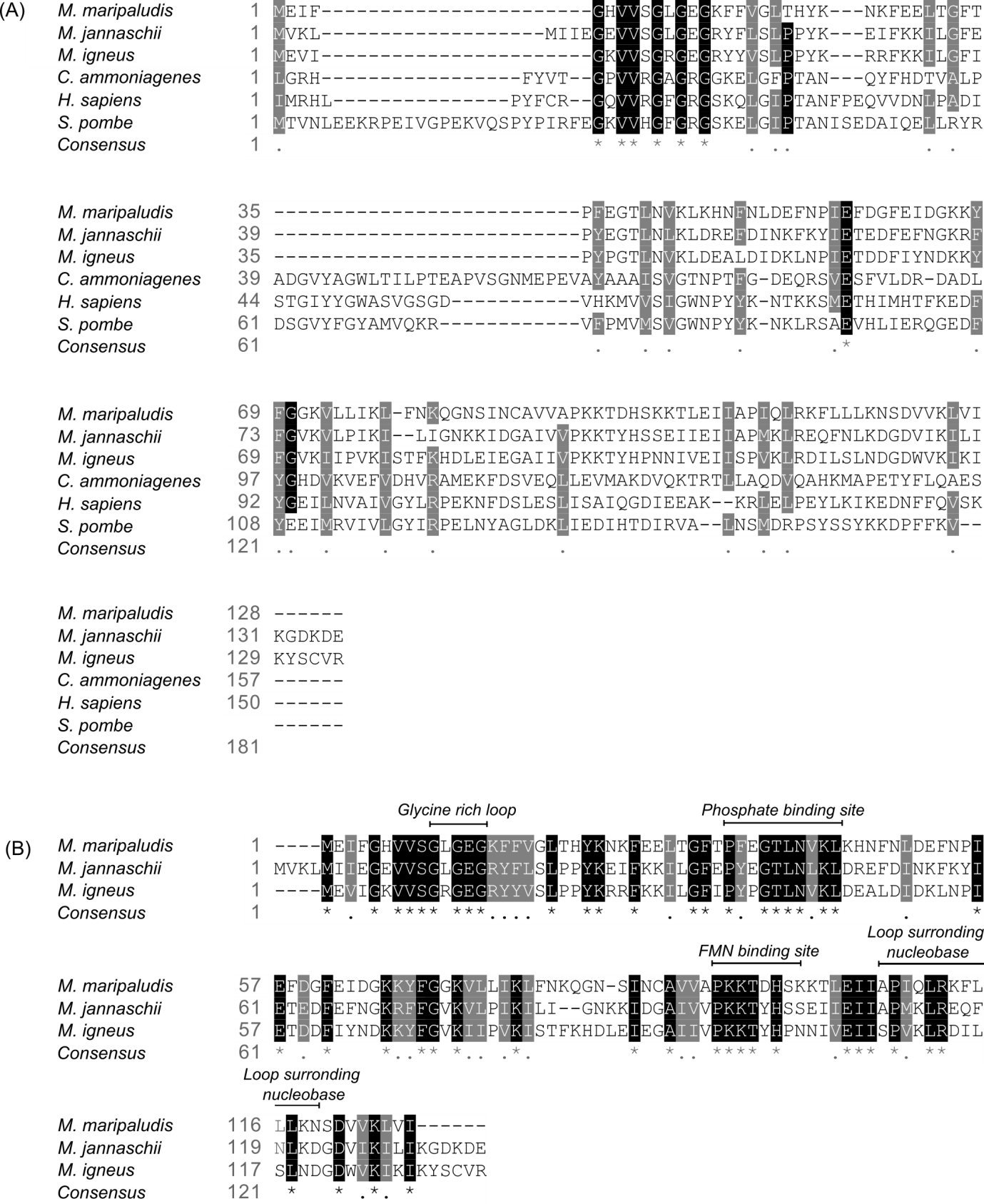

**Figure S1:** **Sequence alignment of archaeal and non-archaeal riboflavin kinases (corresponding to Figure 1 in the manuscript) (A)** Multiple sequence alignment of archaeal and non-archaeal riboflavin kinases shows low overall sequence similarity. Interestingly, some residues in the N-terminal appear to be conserved. **(B)** Multiple sequence alignment (a representative set of three archaeal riboflavin kinases is shown here) shows the well-conserved flavin binding site, nucleobase binding region, and phosphate binding motifs.

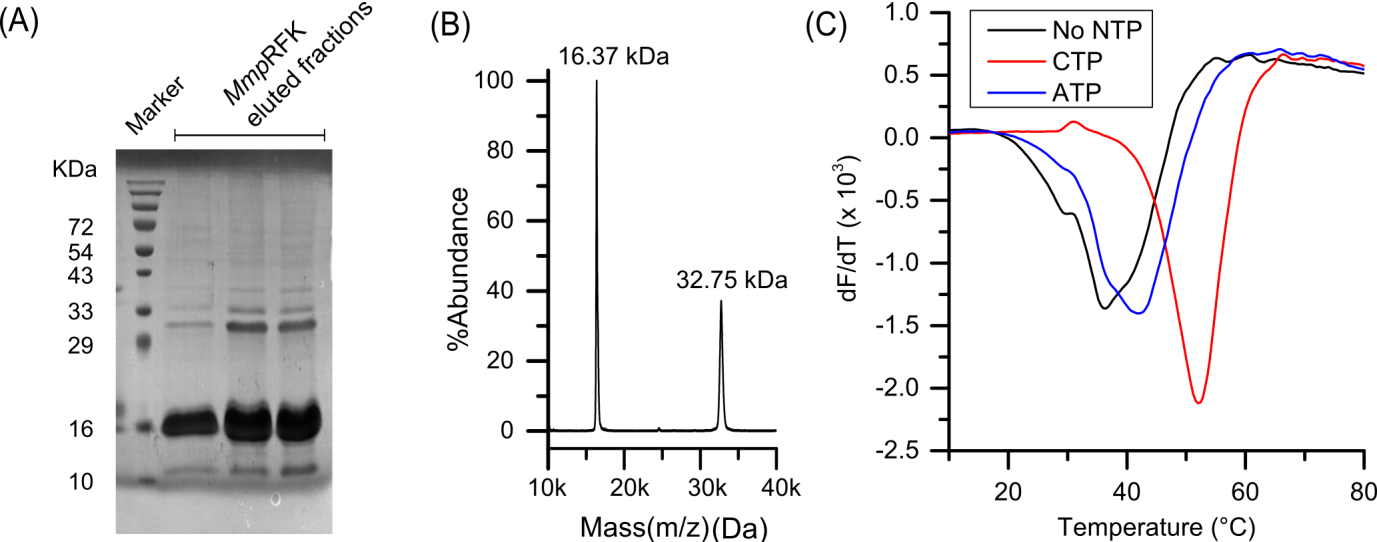

**Figure S2: Protein expression and purification of *Mmp*RibK** **(A)** SDS-PAGE gel electrophoresis showing the 16 kDa band of MmpRibK **(B)** Matrix Assisted Laser Desorption Ionization (MALDI) mass spectrometry data confirms the molecular weight of the purified MmpRibK protein **(C)** Thermal shift assay of *Mmp*RibK with CTP shows that the presence of CTP stabilizes the *Mmp*RibK. When ATP is added to *Mmp*RibK, it does not appear to bind with the protein

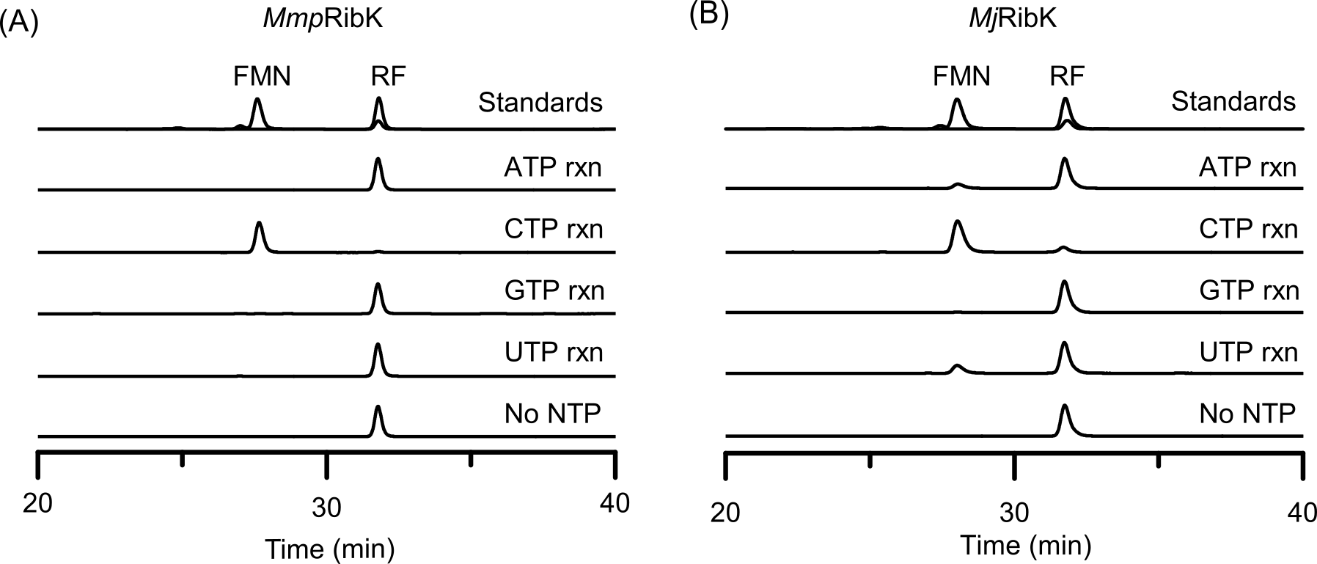

**Figure S3:** **Testing for utilization of alternate nucleotide triphosphate by *Mmp*RibK and *Mj*RibK** **(A)** *Mmp*RibK reactions with riboflavin (RF) and different NTPs shows that it is very specific for CTP for formation of flavin mononucleotide (FMN). **(B)** *Mj*RibK reaction with different NTPs shows that CTP is the preferred substrate. It also gives some conversion with ATP and UTP as also previously noted.^[1]^

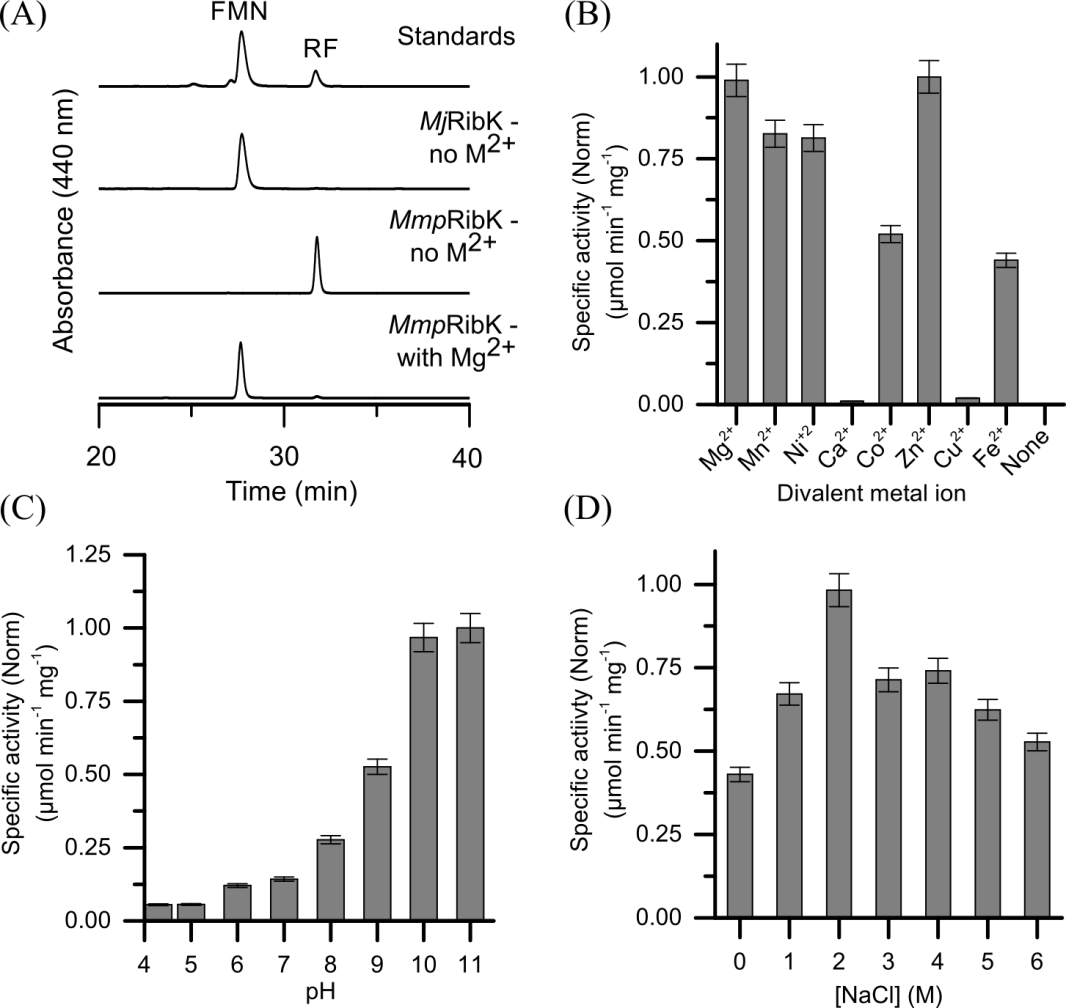

**Figure S4:** Characterization of the biochemical properties of MmpRibK **(A)** *Mmp*RibK and *Mj*RibK reactions with and without metal suggest that the activity of *Mmp*RibK is dependent on metal. *Mj*RibK shows metal independent activity as previously reported (Mashhadi REF).^[1]^ **(B)** Divalent metal ion-dependent activity shows the highest activity of *Mmp*RibK is with Mg^2+^ and Zn^2+^ ions. **(C)** The activity of *Mmp*RibK increases with increasing pH **(D)** *Mmp*RibK shows highest activity at 2M NaCl and can tolerate even saturating concentrations of NaCl.

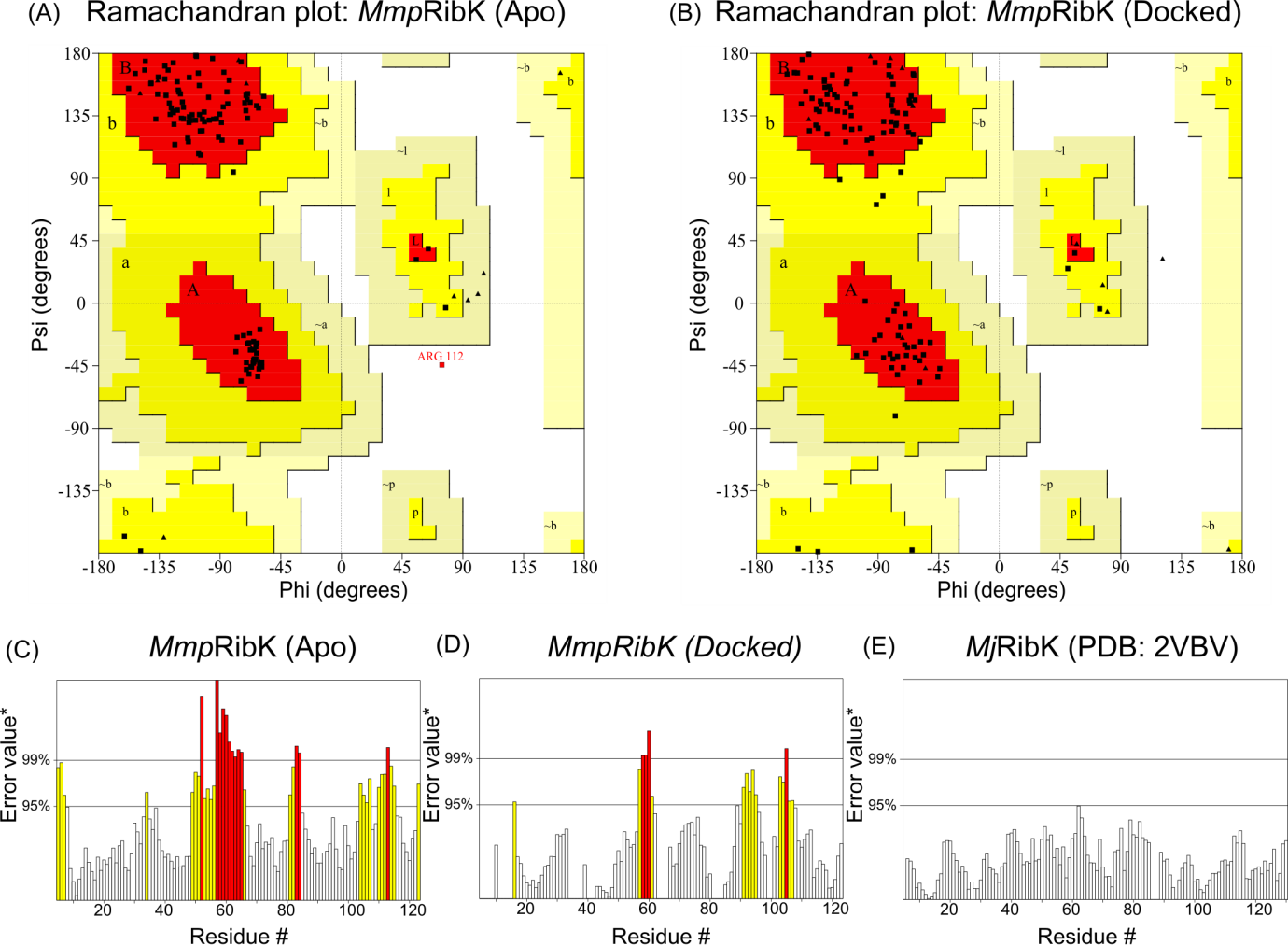

**Figure S5:** **Evaluation of the modeled *Mmp*RibK structure with Ramachandran analysis and ERRAT analysis.** Ramachandran plot of *Mmp*RibK shows the allowed phi-psi dihedral angles of the protein backbone. The red regions are favorable region, and yellow are the allowed regions. The cream regions are generously allowed regions, and white is disallowed regions **(A)** Ramachandran plot of modeled *Mmp*RibK structure in the apo form, which shows that most of the residues (black squares) are in allowed regions. Only one residue, Arg112 (red square), is in the disallowed region. **(B)** Ramachandran plot of *Mmp*RibK docked with CTP and riboflavin. The plot indicating that most of the residues are in the allowed region. Arg112, which was in the disallowed region in the apo form, moved to the allowed region in the docked structure. **(C)** Error-values of residues of *Mmp*RibK apo form using ERRAT analysis indicated the individual quality factors of some residues is poor. **(D)** The corresponding error-values of residues of *Mmp*RibK docked with CTP and riboflavin improve to a large extent **(E)** ERRAT Error values of residues of *Mj*RibK for comparison with the modeled *Mmp*RibK.

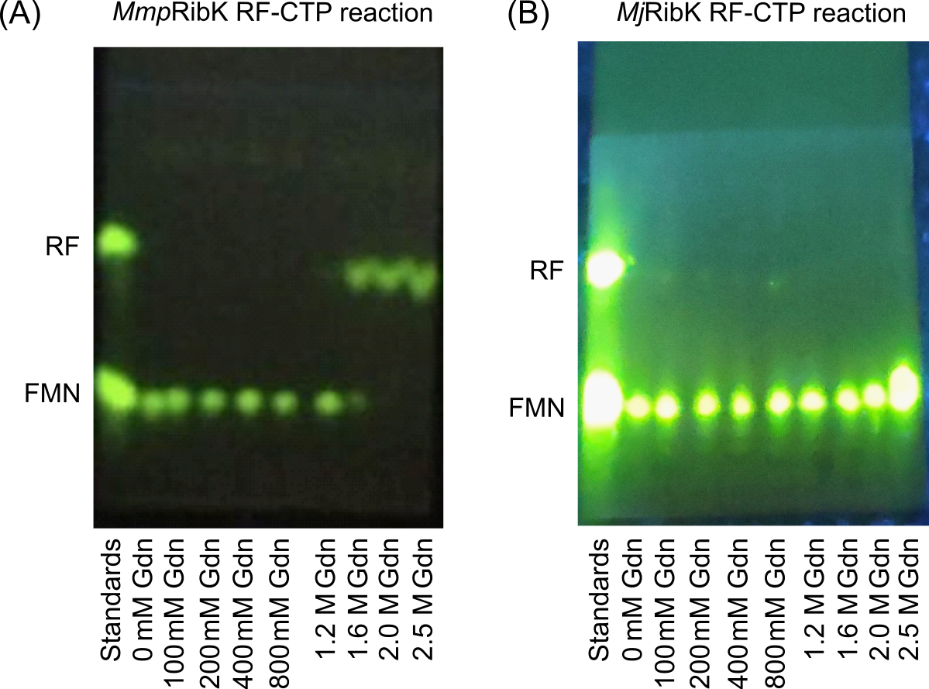

**Figure S6:** **Chemical denaturation of *Mmp*RibK and *Mj*RibK with guanidinium hydrochloride (Gdn-HCl).** **(A)** Activity measurement of *Mmp*RibK in the presence of different concentrations of Gdn-HCl varying from 0 mM to 2.5 M. Enzyme is stable up to 1.2 M Gdn-HCl concentration. After that, it loses its activity. **(B)** Activity measurement of *Mj*RibK in the presence of different concentrations of Gdn-HCl varying from 0 mM to 2.5 M. Enzyme is stable even up to 2.5 M Gdn-HCl concentration and is functional.

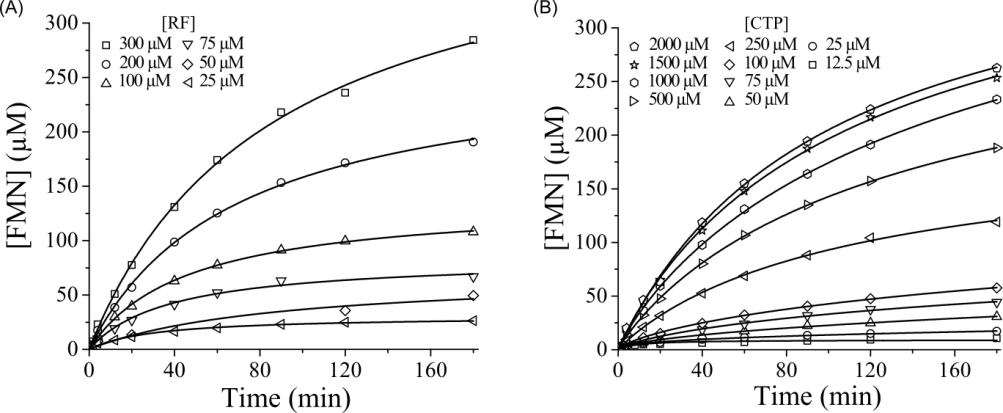

**Figure S7:** **Steady-state kinetics of MmpRibK with substrates riboflavin and CTP: (A)** Initial rates of
product (FMN) formation is measured and plotted at different riboflavin concentrations with a saturating concentration of CTP **(B)** Initial rates of FMN formation is measured and plotted at variable concentrations of CTP with a saturating concentration of riboflavin

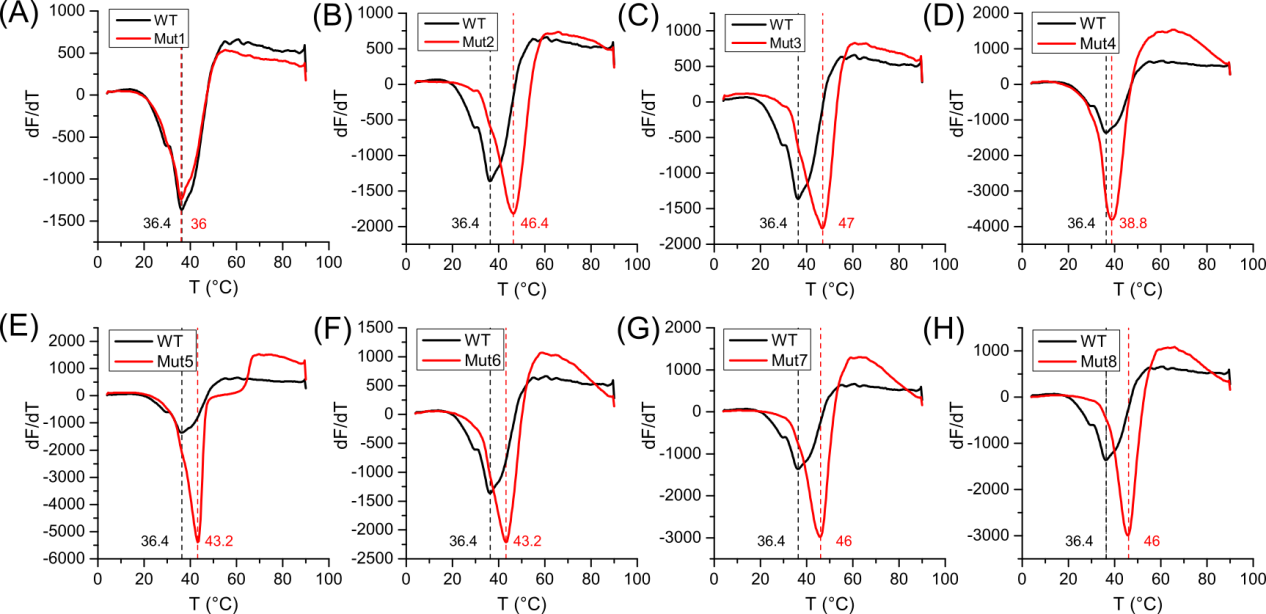

**Figure S8:** **Thermal shift assay of wild-type (WT) *Mmp*RibK and mutants**. We determined the melting temperatures of the wild-type and mutant *Mmp*RibK enzymes, which gives the extent of thermostability introduced by the mutations. **(A)- (E)** The black curves are the melting curves of wild-type, and red curves are the melting curves of mutants. The respective melting temperatures of wild-type and mutants are noted adjacent to their melting curves in black (wild-type) and red (mutants) letters, and the vertical black and red dotted lines indicate these numbers on the x-axis. The shifts of melting temperatures of mutants from the wild-type indicate the extent of thermostabilization.

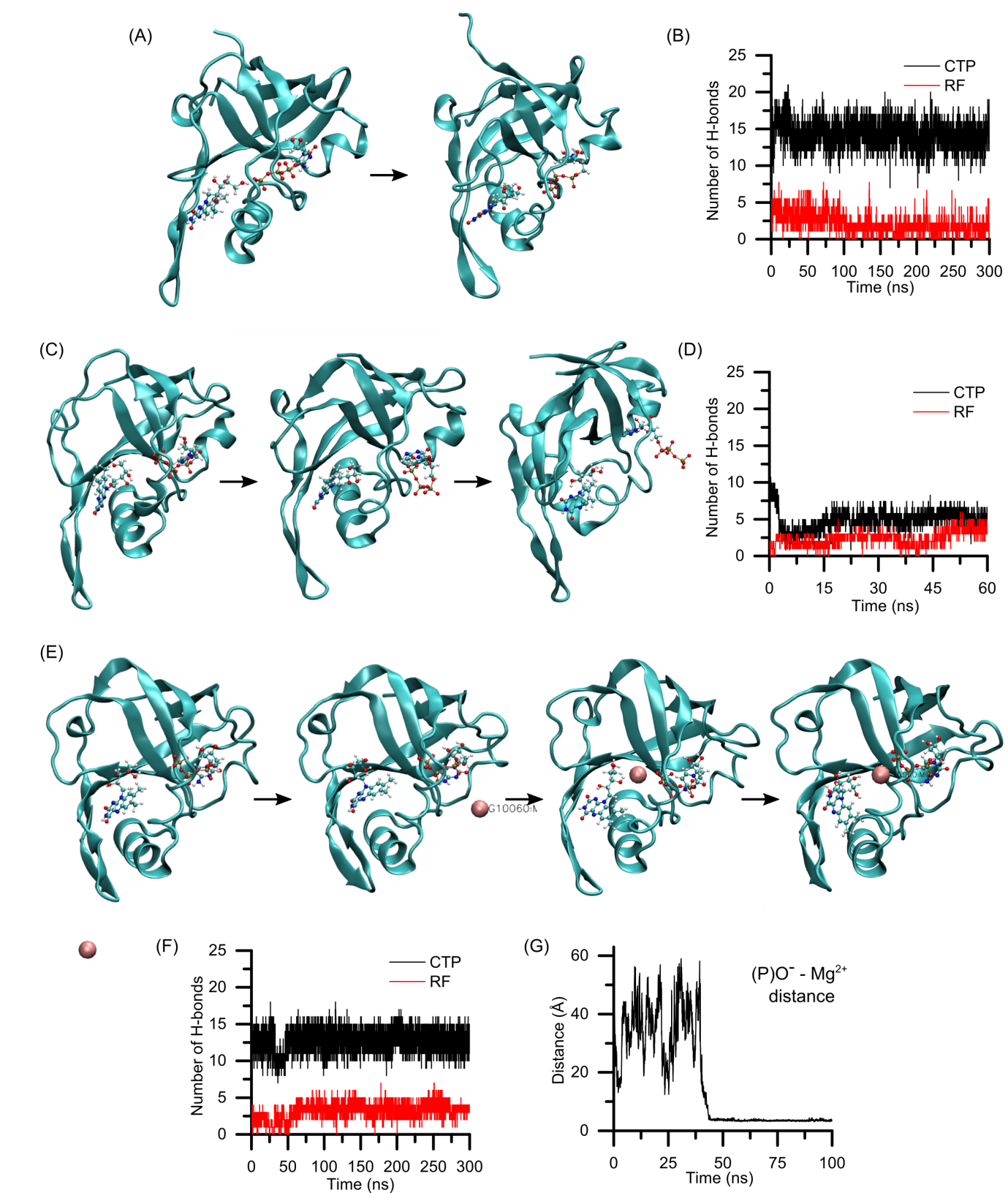

**Figure S9: Molecular dynamics (MD) simulation showing CTP-riboflavin (RF) stability in *Mj*RibK and *Mmp*RibK in the absence and presence of Mg^2+^.** The solid sphere represents the Mg^2+^ ion. Ball and stick model represents the CTP and RF. **(A)** Snapshots of the MD simulation data of the *Mj*RibK active site with CTP and RF in the absence of Mg^2+^. **(B)** The number of hydrogen bonds (H-bonds) between *Mj*RibK and its substrates in absence of Mg^2+^- CTP (black colour) and RF (red colour) **(C)** Snapshots of the MD simulation data of the *Mmp*RibK active site with CTP and RF in the absence of Mg^2+^. The CTP is unable to bind in the active site and comes out **(D)** The number of hydrogen bonds between *Mmp*RibK and CTP (black colour) and RF (red colour) in absence of Mg^2+^. **(E)** Snapshots of the MD simulation data of the *Mmp*RibK active site with CTP and RF in the presence of Mg^2+^. The CTP is now able to bind in the active site and engages with the Mg^2+^ ion **(F)** The number of hydrogen bonds between *Mmp*RibK and CTP (black colour) and RF (red colour) in presence of Mg^2+^. **(G)** Mg^+2^ and phosphate oxygen distance calculated over a 100 ns long molecular dynamic trajectory.

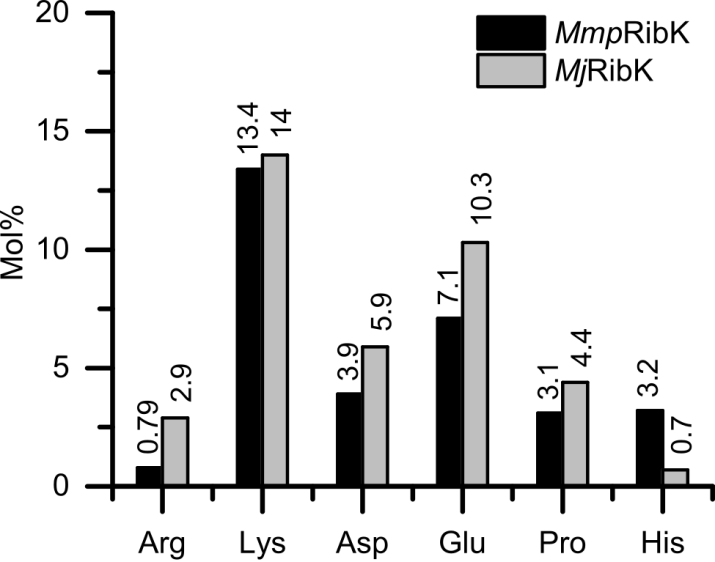

**Figure S10:** Comparison of salt-bridge forming amino acid residues (Arg, Lys, Asp, Glu, His) and rigidifying amino acid residues (Pro) that constitute the sequences of *Mmp*RibK and *Mj*RibK.

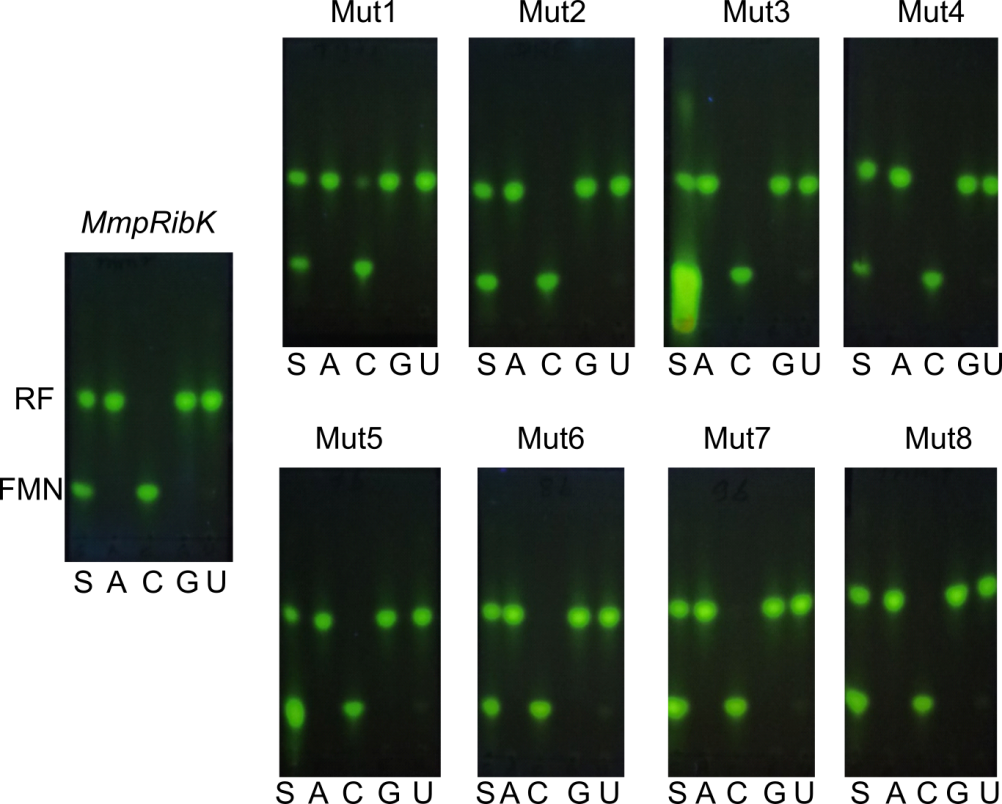

**Figure S11:** **TLC showing the reactions of wild-type *Mmp*RibK and mutants with different NTPs.** S is the standards for riboflavin (RF, top spot) and flavin mononucleotide (FM, bottom spot). A, C, G, and U represents the RibK reactions with riboflavin and nucleotide triphosphates ATP, CTP, GTP and UTP respectively. Wild-type and all the mutants show activity only with CTP, indicating that the mutations have no effect on the nucleotide specificity of *Mmp*RibK.

**Table S1: Top 25 hits from the BLAST analysis with the *Mj*RibK protein sequence**

| # | **Organism** | **Accession no** | **Per. Ident.** | **E-Value** | **Total Score** | **Type** |
| --- | --- | --- | --- | --- | --- | --- |
| 1 | *Methanocaldococcus jannaschii* | 2P3M_A | 100 | 2.52E-88 | 262 | Thermophilic |
| 2 | *Methanocaldococcus jannaschii DSM 2661* | 2OYN_A | 99.265 | 3.30E-88 | 262 | Thermophilic |
| 3 | *Methanocaldococcus sp. FS406-22* | WP_012980051.1 | 89.394 | 3.26E-77 | 234 | Thermophilic |
| 4 | *Methanocaldococcus bathoardescens* | WP_048201574.1 | 87.121 | 1.52E-74 | 227 | Thermophilic |
| 5 | *Methanocaldococcus fervens* | WP_015791162.1 | 85.606 | 1.64E-74 | 227 | Thermophilic |
| 6 | *Methanocaldococcus vulcanius* | WP_015732574.1 | 67.669 | 9.81E-54 | 175 | Thermophilic |
| 7 | *Methanocaldococcus villosus* | WP_004591531.1 | 59.524 | 1.94E-47 | 159 | Thermophilic |
| 8 | *Methanotorris igneus* | WP_013799911.1 | 59.69 | 1.43E-44 | 151 | Thermophilic |
| 9 | *Methanotorris formicicus* | WP_007043845.1 | 59.69 | 2.84E-44 | 150 | Thermophilic |
| 10 | *Methanocaldococcus infernus* | WP_083771775.1 | 59.836 | 1.10E-42 | 147 | Thermophilic |
| 11 | *Methanothermococcus okinawensis* | WP_013866684.1 | 53.968 | 5.79E-42 | 145 | Thermophilic |
| 12 | *Methanothermococcus thermolithotrophicus* | WP_018154369.1 | 56.349 | 2.20E-41 | 143 | Thermophilic |
| 13 | *Methanococcus aeolicus* | WP_011973638.1 | 52.5 | 5.90E-38 | 134 | Mesophilic |
| 14 | *Methanofervidicoccus abyssi* | WP_131007122.1 | 47.619 | 2.26E-37 | 133 | Thermophilic |
| 15 | *Methanococcus maripaludis DSM 2067* | AVB76310.1 | 52.308 | 5.99E-36 | 129 | Mesophilic |
| 16 | *Methanofervidicoccus sp. A16* | WP_148120329.1 | 46.457 | 4.24E-35 | 127 | Thermophilic |
| 17 | *Methanococcus maripaludis C5* | ABO35789.1 | 52.273 | 7.15E-35 | 127 | Mesophilic |
| 18 | *Methanococcus maripaludis C6* | WP_012193477.1 | 53.543 | 2.66E-34 | 125 | Mesophilic |
| 19 | *Methanococcus maripaludis C7* | WP_011977559.1 | 51.181 | 3.03E-34 | 125 | Mesophilic |
| 20 | *Methanococcus maripaludis OS7* | WP_119720556.1 | 51.163 | 2.72E-33 | 123 | Mesophilic |
| 21 | *Methanococcus vannielii* | WP_012065999.1 | 52.381 | 5.99E-33 | 122 | Mesophilic |
| 22 | *Methanococcus maripaludis KA1* | WP_146777918.1 | 50.388 | 6.31E-33 | 122 | Mesophilic |
| 23 | *Methanococcus maripaludis X1* | WP_013998637.1 | 50.388 | 1.70E-32 | 120 | Mesophilic |
| 24 | *Methanococcus maripaludis S2* | WP_011170128.1 | 50.388 | 5.98E-32 | 119 | Mesophilic |
| 25 | *Methanococcus voltae* | WP_013180855.1 | 45.802 | 2.21E-29 | 113 | Mesophilic |

**Table S2: Ramachandran plot statistics**

| Plot statistics | *Mmp*RibK (Apo) | | *Mmp*RibK (Docked) | | *Mj*RibK | |
| --- | --- | --- | --- | --- | --- | --- |
| Residues in most favoured regions [A,B,L] | 104 | 95.40% | 99 | 90.80% | 97 | 85.10% |
| Residues in additional allowed regions [a,b,l,p] | 4 | 3.70% | 10 | 9.20% | 17 | 14.90% |
| Residues in generously allowed regions [~a,~b,~l,~p] | 0 | 0.00% | 0 | 0.00% | 0 | 0.00% |
| Residues in disallowed regions | 1 | 0.90% | 0 | 0.00% | 0 | 0.00% |
| Number of non-glycine and non-proline residues | 109 | 100.00% | 109 | 100.00% | 114 | 100.00% |
| Number of end-residues (excl. Gly and Pro) | 2 |  | 4 |  | 2 |  |
| Number of glycine residues (shown as triangles) | 12 |  | 12 |  | 12 |  |
| Number of proline residues | 4 |  | 4 |  | 6 |  |
| Total number of residues | 127 |  | 129 |  | 134 |  |

**Table S3: Comparison of mol% of amino acid composition between *Mmp*RibK and *Mj*RibK**

| **Amino acid** | ***Mmp*RibK** | ***Mj*RibK** | **Amino acid** | ***Mmp*RibK** | ***Mj*RibK** |
| --- | --- | --- | --- | --- | --- |
| Ala | 2.36 | 1.47 | Met | 0.79 | 2.21 |
| Cys | 0.79 | 0.00 | Asn | 7.09 | 3.68 |
| Asp | 3.94 | 5.88 | Pro | 3.15 | 4.41 |
| Glu | 7.09 | 10.29 | Gln | 1.57 | 0.74 |
| Phe | 10.24 | 7.35 | Arg | 0.79 | 2.94 |
| Gly | 9.45 | 8.82 | Ser | 3.15 | 2.94 |
| His | 3.15 | 0.74 | Thr | 4.72 | 2.21 |
| Ile | 7.09 | 13.24 | Val | 7.87 | 5.88 |
| Lys | 13.39 | 13.97 | Trp | 0.00 | 0.00 |
| Leu | 11.81 | 9.56 | Tyr | 1.57 | 3.68 |

**Table S4: List of the primers used in the cloning**

| # | Primer name | Sequence |
| --- | --- | --- |
| 1 | *Mj*RibK_F1 | 5'-TTGGTGAAATTGATGATTATTG-3' |
| 2 | *Mj*RibK_R1 | 5'-TTATTCATCTTTATCTCCC-3' |
| 3 | *Mj*RibK_F2 | 5'-GCCGCGCGGCAGCCATATG GTGAAATTGATGATTATTG-3' |
| 4 | *Mj*RibK_R2 | 5'-CGACGGAGCTCGAATTCGGATCCTTATTCATCTTTATCTCCC-3' |
| 5 | *Mmp*RibK_F1 | 5'-ATGGAAATTTTTGGGCATG-3' |
| 6 | *Mmp*RibK_R1 | 5'-TTAAATTACGAGTTTTAC-3' |
| 7 | *Mmp*RibK_F2 | 5'-GCCGCGCGGCAGCCATATGGAAATTTTTGGGCATG-3' |
| 8 | *Mmp*RibK_R2 | 5'-CGACGGAGCTCGAATTCGGATCCTTAAATTACGAGTTTTAC-3' |
| 9 | *Mmp*RibK_T34E_F1 | 5'-GAATTAACCGGATTTGAACCGTTTGAAGGAACA-3' |
| 10 | *Mmp*RibK_S2_DRE_F1 | 5'-AGGAACATTGAACGTAAAATTAGATAGAGAATTCAATTTAGATGAATTTAATCC-3' |
| 11 | *Mmp*RibK_N119D_R1 | 5'-TTTTACAACATCTGAATCCTTTAATAATAAAAA-3' |
| 12 | *Mmp*RibK_TH(21-22)PP_F1 | 5'-AAGTTTTTTGTTGGATTACCTCCTTACAAAAATAAATTTGAA-3' |
| 13 | T7 promoter_F | 5'-TAATACGACTCACTATAGGG-3' |
| 14 | T7 terminator_R | 5'-GCTAGTTATTGCTCAGCGG-3' |

**Table S5: List of the plasmids constructed in the study**

| **#** | **wild-type and mutants** | **Plasmid name** |
| --- | --- | --- |
| 1 | *Mj*RibK | pYK1701 |
| 2 | *Mmp*RibK | *pYK1801* |
| 3 | *Mut1* | *pYK1908* |
| 4 | *Mut2* | *pYK1902* |
| 5 | *Mut3* | *pYK1910* |
| 6 | *Mut4* | *pYK1909* |
| 7 | *Mut5* | *pYK1911* |
| 8 | *Mut6* | *pYK1912* |
| 9 | *Mut7* | *pYK1913* |
| 10 | *Mut8* | *pYK1914* |

**Supplementary methods**

**Effect of NaCl concentration on *Mmp*RibK activity:** The enzyme activity was assayed in varying salt conditions, ranging from 50 mM to saturating NaCl concentration (solid NaCl is weighed and added directly to the reaction mixture for higher saturating concentrations of NaCl). 100 µM of riboflavin and 4 mM CTP is used in this experiment, and the reaction was analyzed by TLC and HPLC as described in the method section.

**Test for alternate nucleotides as co-substrate for riboflavin phosphorylation:** The enzyme activity of *Mmp*RibK and its mutants were tested with alternate co-substrate nucleotides ATP, GTP, and UTP, both in the absence and presence of NaCl, and analyzed by TLC and HPLC as described in the method section.

**Metal ion dependence of *Mmp*RibK*:*** The enzyme assay as previously described was performed in the presence of 10 mM of MgCl_2_.6H_2_O, MnCl_2_.4H_2_O, NiCl_2_.6H_2_O, CaCl_2_.H_2_O, CoCl_2_.6H_2_O, ZnCl_2_, CuCl_2_.2H_2_O, FeSO_4_.7H_2_O (freshly prepared), and the reaction was analysed using TLC and HPLC.

**Temperature dependence of *Mmp*RibK:** The optimal reaction temperature for *Mmp*RibK activity with riboflavin, CTP, and Mg^2+^ was determined by conducting the assay in the temperature range 15°C-70°C, and analyzing it as described previously.

**pH dependence of *Mmp*RibK*:*** The optimal pH for the enzyme activity of *Mmp*RibK with riboflavin, CTP, and Mg^2+^ was determined by conducting the assay in the pH range of 4.3-11, and analyzing it as described previously. The following buffers were used to achieve the different pH conditions - sodium acetate buffer for pH 4.3 and 5.0, MES buffer for pH 6.0, HEPES for pH 7.0, and Tris-HCl buffer for pH 8.0, 9.0, 10.0, and 11.0.

**Chemical denaturation of *Mmp*RibK and *Mj*RibK:** The activities of *Mj*RibK and *Mmp*RibK were measured in the presence of chemical denaturant guanidinium hydrochloride (Gdn-HCl), to check the difference in the stability of these thermophilic-mesophilic pair of the enzyme. Variable concentrations of Gdn-HCl (0 mM – 2.5 M) were added to the reaction mixture containing 100 µM of riboflavin, 100 mM Tris-HCl pH 8.0, 10 mM MgCl_2_, 1 µM of the enzyme. These are kept at room temperature for one hour. After one-hour, samples of *Mmp*RibK were incubated at 37°C, and those of *Mj*RibK were set at 70°C for 20 min. The reactions were initiated by adding 2 mM of CTP and quenched with acetic acid after 15 min. The reaction product FMN was analysed on TLC (Fig S5)

**Compuational Method and Details**

**Modelling of Protein:** The 3-dimensional structure of *Mmp*RibK is not available, however, its amino acid sequence is known. Therefore, first we modeled the 3-dimensional structure of the protein from the sequence and then performed molecular dynamic simulation at temperatures, 27°C and 72°C, to understand the stability.

Getting an accurate model of an unknown protein is very challenging. However, if the unknown protein sequence is very similar to a known protein (whose structure is already known), the modeled protein will likely have similarity to the template protein and the predicted conformation will be the most probable conformation. Here, in the present work we have used the MODELLER module to model the protein, which works on the principle of the homology modelling, where we provide the alignment of sequence to be modeled with the known structures.^[2]^ Then, MODELLER uses a spatial restraints objective function to optimize the modeled protein. *Mmp*RibK is member of kinase family, therefore its structure is very similar (50%) to the *Mj*RibK and since the crystal of *Mj*RibK is known (PDB ID: 2VBV ), MODELLER used *Mj*RibK as a template for creating the *Mmp*RibK structure. The structure of modeled protein (*Mmp*RibK) superimposed over the template protein (*Mj*RibK) is shown in Figure 2C. Since there is 50 % similarity between the proteins, the 3-D structure is very similar (RMSD 1.2 Å).

After, modeling we performed a micro-second simulation to see its stability (see simulation detail)

**Simulation Detail:** After modeling the *Mmp*RibK, we performed molecular dynamic simulation. The AMBER99SB force field and initial conformation of protein was generated by GROMACS.^[3,4]^ The protein was kept in cubic box of dimension of 70 Å and solvated with ~2000 TIP3P water molecules. Thereafter, 150 mM of MgCl_2_ was added to neutralize the system, followed by the energy minimization using steepest descent method.^[5]^ After the energy minimization, system was heated to 27°C in 4 ns in NVT condition by using the Berendson thermostat with coupling constant of 0.2 ps. Subsequent to that we have performed 5 ns NPT simulation using Berendsen thermostat and barostat with coupling constant of 0.2 ps.^[6]^ The NPT simulation was performed to adjust the box size of the system.

Finally, we have performed 1 μs NVT simulation using the Nose-Hoover thermostat.^[7]^ During the simulation, all bonds were constrained using LINCS algorithm.20 Particle Mesh Ewald (PME) method was used for the electrostatics with a 10 Å cut-off for the long range interaction. Cut-off for the van der Waals (vdW) interaction was kept at 10 Å. Simulations was performed with 2 fs time step. All simulation was done by using GROMCS.

**Docking Study**: *Mmp*RibK converts RF to FMN by utilizing a CTP cofactor which is converted to CDP after reaction. However, the binding site of CTP and RF is not known. Therefore, we docked the CTP and RF to *Mmp*RibK to make *Mmp*RibK-CTP-RF complex. A gross idea of CTP and RF binding site is guessed from its homologous protein *Mj*RibK. Residue VAL7, SER9, GLY10, LEU11, GLY12, GLU13, GLY14, LYS15, PHE36, GLY38, THR39, LEU40, ASN41, ILE06, ALA107, ILE109, GLN110, LEU111, ARG112, LEU117, LYS118, ASN119 was used for docking of CTP and PHE17, TYR23, TYR68, PHE69, GLY70, VAL91, ALA92, PRO93 and GLU104 used for RF docking. We used Autodock tool for the docking and first RF was docked and then CTP.^[8,9]^ The grid size for docking at above mention active side was 15, 15 and 27 respectively for x, y and z direction.

After Docking the CTP and RF, we performed the molecular dynamic simulation. Here, also we used AMBER99SB forcefiled for protein. However, to create the amber forcefield of CTP and RF we used antechamber module of Ambertool.^[10]^ First, we optimized the CTP quantum mechanically using HF/6-31G* basis set in GAUSSIAN03 software and then restricted electrostatic potential (resp) charges on the atoms of CTP and RF were calculated.^[11]^ The topology and co-ordinates generated from AmberTools were converted into GROMACS format using a perl program amb2gmx.pl. We put the CTP-RF bound protein in a cubic box of length 70 Å. To study the effect of metal on CTP and RF stability we performed two set of simulation of *Mmp*RibK-CTP-RF complex in (i) absence and (ii) in presence of Mg^2+^. All other simulations details are same as mentioned above. The only difference is that we performed 300 ns long production at 27°C and at 72°C using Nose-Hoover thermostat. All the analysis is done over a production run. We used GROMACS module for the analysis.

**Creating the mutants:** The *Mmp*RibK, a mesophilic protein, is operational at room temperature (27°C) whereas its homolog *Mj*RibK (a thermophilic protein) operates at 72°C temperature. To convert the *Mmp*RibK to a thermophilic protein we created the 8 mutant of *Mmp*RibK (see table 3 of MS), based on the salt bridges and non-conserved proline residues of the protein. To create the mutant we used the Swapaa module of Chimera.^[12]^ All the mutations were made in the CTP and RF bound complex. After mutation, each system was simulated for 300 ns in 150 mM MgCl_2_ solution. All the simulation details are same as mentioned above and the analysis was performed over a 300 ns long simulation.
